# Constitutive macropinocytosis sustains the inflammatory phenotype of senescent cells

**DOI:** 10.64898/2026.07.12.738086

**Authors:** Kamila Kozik, Nurhanani Razali, Keiko Kono

## Abstract

**Summary:** Senescent cells upregulate constitutive macropinocytosis that is required for inflammatory SASP production through PAK1 and TGFβ signaling, revealing a novel role for macropinocytosis in the senescence associated secretory phenotype.

Cellular senescence is a stable cell cycle arrest characterized by extensive metabolic remodeling and a proinflammatory secretome known as senescence-associated secretory phenotype (SASP). While transient SASP supports tissue repair and wound healing, its persistent activation drives chronic inflammation and age-related pathology. Despite the complex regulation of SASP, the contribution of vesicle trafficking and endocytic pathways to its control remains poorly defined. Here, we identify macropinocytosis as a constitutively active endocytic pathway in multiple cell lines and senescence subtypes. Using pharmacological and genetic perturbations, we demonstrate that PAK1 is a key regulator of macropinocytosis in senescent cells. Moreover, PAK1-dependent macropinocytosis regulates production of inflammatory SASP factors, such as IL6 and CCL2, through TGFβ signaling. Our findings define senescence-associated macropinocytosis as a mechanistic link between endocytosis and the proinflammatory phenotype of senescent cells, establishing PAK1 and macropinocytosis as potential therapeutic targets for alleviating the deleterious effects of senescence in aging and cancer.

## Introduction

Cellular senescence is a stable cell cycle arrest that emerges in response to various stressors, including telomere attrition, oncogenic signaling, chemotherapeutic exposure, DNA damage response and plasma membrane damage^1–8^. Cellular senescence functions primarily as a tumor-suppressive and damage-limiting program that arrests the division of damaged cells; however, sustained accumulation of senescent cells is deleterious, as it promotes chronic inflammation, disrupts tissue homeostasis, regenerative capacity, and contributes to multiple age-related disorders^9–13^. These pathological consequences arise largely from the senescence-associated secretory phenotype (SASP), a complex and heterogeneous mixture of proinflammatory cytokines, chemokines, matrix-remodeling enzymes, extracellular vesicles and growth factors that varies depending on the senescence trigger, cell type, and stage of senescence^14–18^. Consequently, the upstream mechanisms that regulate SASP are only partially understood. Strategies aimed at removing or preventing cellular senescence, including pharmacological clearance, senescence reprogramming, or modulation of the proinflammatory SASP, hold promise for promoting healthy aging^19–25^.

Senescent cells undergo profound morphological remodeling^26–32^. These structural alterations are accompanied by changes in membrane trafficking pathways, including endocytosis, which regulates nutrient uptake, receptor availability, and downstream signaling^33–36^. In senescent cells, clathrin- and caveolin-mediated endocytosis are downregulated, linking altered trafficking to the senescence-associated phenotypes^37–40^. Collectively, findings on these two endocytosis pathways suggest a broader reorganization of membrane dynamics that supports metabolic rewiring, inflammatory signaling, and long-term survival in senescent cells. However, the involvement of other endocytic pathways in senescence remains largely unexplored.

Emerging evidence suggests that senescent cells accumulate large cytoplasmic vacuoles, a feature characteristic of macropinocytosis^41,42^. Macropinocytosis is an actin-driven, non-selective endocytic process in which large volumes of extracellular fluid are internalized through macropinosomes^43,44^. Macropinocytosis plays critical roles in immune surveillance in immune cells and nutrient scavenging in tumor microenvironment^45,46^. For example, cancer-associated fibroblasts (CAFs) support tumor metabolism through enhanced secretion of amino acids and nutrients fueled by upregulated macropinocytosis, and Ras-transformed cancer cells rely on macropinocytosis to adapt to metabolic stress^47,48^. While extensively studied in cancer, immune cells, and nutrient-deprived systems, macropinocytosis has been largely unexplored in senescent cells.

One of the key regulators of macropinocytosis is PAK1 (p21-activated kinase 1), a serine/threonine kinase activated downstream of Rac1 and Cdc42, often via PI3K signaling^49–53^. PAK1 coordinates actin cytoskeletal remodeling by phosphorylating targets involved in membrane ruffling and lamellipodia formation, processes essential for macropinosome formation^54^. It promotes the closure of membrane ruffles into macropinosomes, thereby enabling efficient uptake of extracellular fluid and solutes^52^. PAK1 activity is also required for the maturation and trafficking of macropinosomes^55^. PAK1 therefore serves as a central effector of macropinocytosis in various physiological and pathological contexts. Additionally, recent evidence positions PAK1 as an emerging aging kinase^56^. Genetic inhibition of PAK1 extends lifespan and delays functional decline in *C. elegans*, in part by promoting the activity of the pro-longevity transcription factor DAF-16/FOXO^56^. These findings suggest that PAK1 may suppress stress resistance and longevity programs in certain contexts. However, its precise role in mammalian senescence remains unclear, and how PAK1 mechanistically regulates the senescence phenotype or SASP is not well understood.

In this study, we show that macropinocytosis is constitutively upregulated in senescent cells across multiple senescence and cell types. We identify PAK1 as a central driver of macropinocytosis in senescent cells, demonstrating that this pathway sustains inflammatory cytokine production via TGFβ signaling and uncovering a novel link between endocytosis and the senescence-associated secretory phenotype. These findings highlight senescence-associated macropinocytosis and PAK1 as a potential therapeutic target for mitigating the detrimental effects of chronic senescence in aging and cancer.

## Results

### Senescent cells upregulate peripheral large vesicle formation associated with extensive membrane ruffling

Senescent cells contain large cytoplasmic vesicles^41,57–60^. However, the molecular mechanisms underlying their formation and persistence in cytoplasm remain incompletely understood. To characterize this phenotype, we investigated cytoplasmic vacuolization across multiple senescence models. We first validated senescence induction in normal human fibroblasts WI-38 and BJ cells, and human osteosarcoma U2OS cells following three distinct senescence triggers: replicative exhaustion, DNA damage, and plasma membrane damage (Fig. 1a). For replicative exhaustion (replicative senescence, Rep-Sen), WI-38 and BJ fibroblasts were repeatedly passaged until they reached a proliferative arrest (population doubling level >50 for WI-38 and >70 for BJ cells). Stress-induced senescence was induced by either the genotoxic drug doxorubicin (250 nM, 24 h) (DNA damage response-dependent senescence, DDR-Sen) or the membrane-damaging agent sodium dodecyl sulfate (SDS, 0.009% for WI-38 and 0.014% for BJ cells, 24 h) (plasma membrane damage-dependent senescence, PMD-Sen) as previously reported^7^. After the treatment, the medium was changed and the cells were maintained in culture for an additional 16 days to allow full establishment of senescence (Fig. 1a). In U2OS cells, senescence was induced using a lower dose of doxorubicin (100 nM for 24 h). Successful induction of senescence in WI-38 cells was confirmed by SA-β-Gal staining (Fig. 1b and c), loss of Lamin B1 and upregulation of p21 and p16 by western blotting (Fig. 1d), and increased mRNA levels of SASP factors *IL6* and *CCL2* by qPCR (Fig. 1e). Additionally, WI-38 senescent cells displayed a significant enlargement of nuclear area, consistent with characteristic morphological remodeling (Fig. 1f). Senescence in BJ and U2OS was confirmed by SA-β-Gal staining (Supplementary Fig. 1a-d), and western blotting (Supplementary Fig. 1e and f). These results confirm successful senescence induction in WI-38, BJ and U2OS cells.

**Figure 1.**
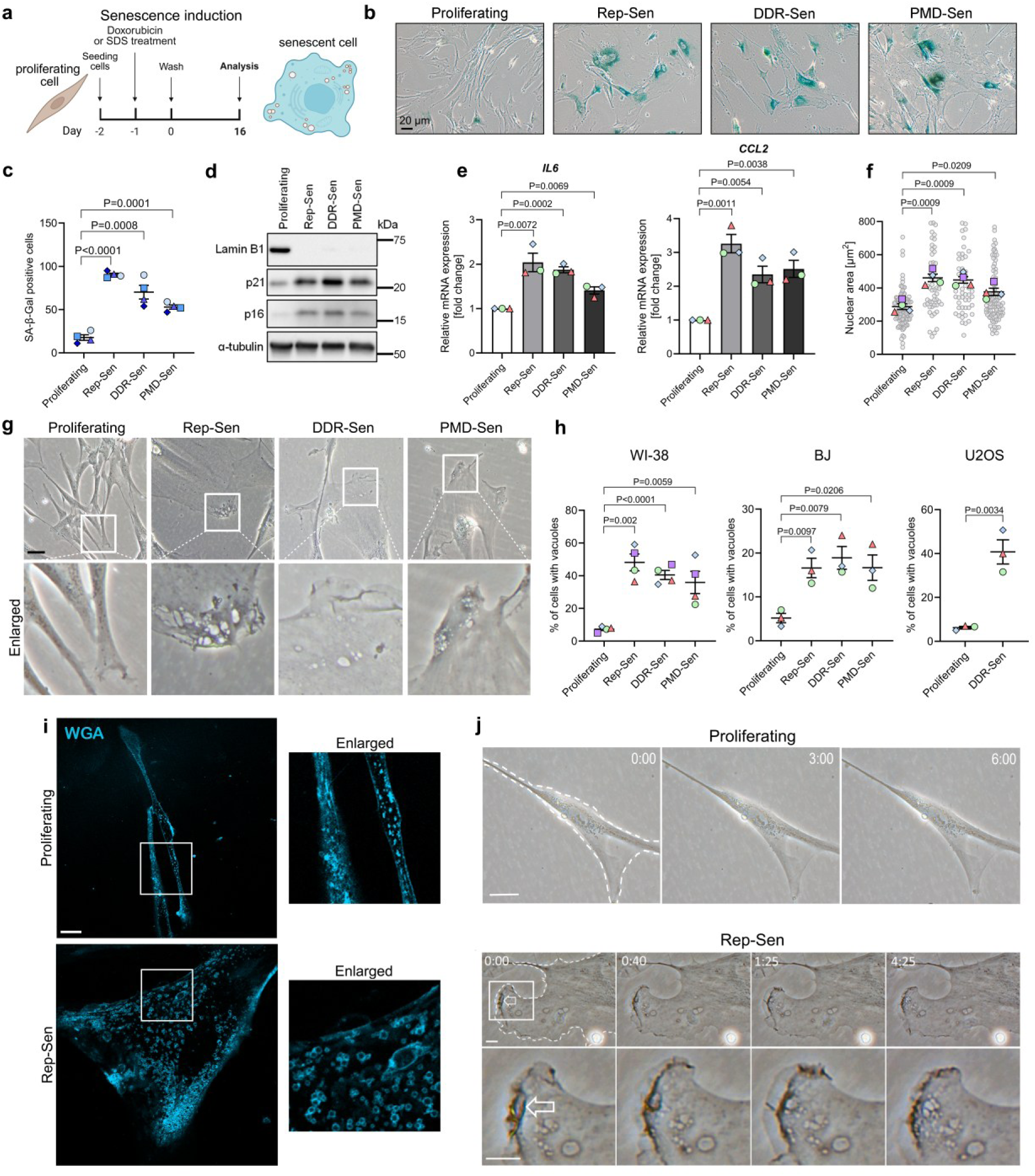
Senescent cells form a high number of peripheral cytoplasmic large vesicles. **(a)** Scheme of senescence induction**. (b)** SA-β-Gal positive cells detected after replication arrest (Rep-Sen) or 16 days after treatment (doxorubicin – DDR-Sen, SDS – PMD-Sen). Scale bar 20 μm. **(c)** Quantification of A. Approximately 100 cells were counted per experiment. n=4. **(d)** Western blotting using lysates of proliferating and senescent WI-38 cells. The lysates were collected 16 days after treatment with SDS or doxorubicin. **(e)** qPCR analysis of *IL6* and *CCL2* gene expression. RNA was isolated on day 16 of senescence. n=3. **(f)** WI-38 cells were incubated with DAPI for 30 min and nuclear area was quantified. n=4. **(g)** Phase-contrast analysis of vacuole number in proliferating and senescent WI-38. Scale bar 10 μm. Arrows mark the phase-bright cytoplasmic vacuoles. **(h)** Quantitative analysis of data shown in g. Approximately 50 cells were counted per experiment. n=4 or 3. **(i)** Staining of membrane-bound vesicles using WGA 488 in WI-38 senescent and proliferating cells. Scale bar 20 μm **(j)** Representative frames from time lapse recordings (Movie 1 and 4) of vesicles formation in proliferating (up, Movie 4) and senescent (down, Movie 1) WI-38 cells. Scale bar 20 and 10 μm, respectively. Time format m:ss.

We next quantified the proportion of cells with large cytoplasmic vesicles in proliferating and senescent populations. Across all senescence-inducing conditions, we observed an accumulation of prominent vacuolar structures visible under bright-field microscopy (Fig. 1g and h, Supplementary Fig. 1g). To verify that these structures represent membrane-bound vesicles rather than optical artifacts, we used Alexa Fluor 488-conjugated wheat germ agglutinin (WGA-488), a lectin that labels N-acetylglucosamine-containing glycoproteins on the plasma membrane and on internalized vesicles^61^. Following a 60-min incubation at 37 °C, senescent WI-38 cells internalized WGA, showing large, membrane-associated vesicles throughout the cytoplasm and confirming large vesicle accumulation in the senescent and not in proliferating cells (Fig. 1i). Notably, most vacuoles in senescent cells were not localized to the perinuclear region, where endolysosomal compartments are typically enriched. Instead, they were observed at the cell periphery near the plasma membrane. To verify the peripheral localization of these vacuoles, we performed live-cell imaging and observed extensive plasma membrane ruffling followed by formation of newly emerging peripheral vesicles in all subtypes of senescent but not proliferating WI-38 cells (Fig. 1j, Supplementary Fig. 1h-i and Movie 1-4).

We next aimed to characterize the identity and composition of the endosomal populations contributing to senescence-associated vacuolization. To this end, we generated U2OS cell lines stably expressing GFP-tagged endosomal markers, including Rab5 (early endosomes), Rab4 (recycling endosomes), Rab20, Rab21, and Rab8 (Rab GTPases associated with macropinosomes), Rab7 and LAMP1 (late endosome/lysosome markers), Phafin2 (a regulator of macropinosome maturation), and a tandem 2×FYVE domain probe (PtdIns3P) using lentiviral transduction to ensure genomic integration and stable expression^62,63^. Following induction of senescence with doxorubicin, we observed a marked increase in the number of large peripheral vesicles positive for FYVE, Phafin2, and Rab5 compared with proliferating cells (Supplementary Fig. 1j). In addition, Rab8- and Rab21-positive compartments displayed altered localization and abundance in senescent cells, with a pronounced increase in Rab8- and Rab21-positive vesicles relative to the proliferative state (Supplementary Fig. 1j).

Collectively, these findings indicate a consistent increase in cytoplasmic vesicles in replicative and premature senescent cells, across multiple cell types including WI-38 and BJ fibroblasts, and U2OS osteosarcoma cells. Moreover, accumulation of cytoplasmic vesicles at the cell periphery suggests that they may arise through processes linked to plasma membrane remodeling or internalization. Together, these changes indicate senescence-associated remodeling of early endosomal and macropinosome-related compartments.

### Large vesicle formation and trafficking are rapid, spatially regulated, and cytoskeleton-dependent in senescent cells

To quantitatively assess large vesicle formation in senescent cells, we used holotomographic microscopy on proliferating and senescent WI-38 cells. This label-free phase imaging reconstructs three-dimensional refractive index maps of cells, allowing high-resolution visualization of dynamic subcellular membranes^64^. Using this approach, we observed membrane ruffling (Fig. 2a) and rapid *de novo* formation of vacuolar structures which matured into spherical vesicles and moved centripetally toward the cell interior in senescent cells (Fig. 2a-c and Movie 5-7). Timelapse holotomography confirmed phase-contrast observations that certain vacuoles in senescent cells form within seconds (Fig. 2c and Movie 5). Immediately after their appearance at regions of intense ruffling, these nascent vesicles had elongated or tubular morphologies (Fig. 2b). These nascent vesicles subsequently fused to form large (>2 μm²), round vesicles (Fig. 2c). Quantitative analysis showed a significant increase in both the number and size (area and volume) of vacuoles in senescent WI-38 cells compared with their proliferating counterparts (Fig. 2d and e).

**Figure 2.**
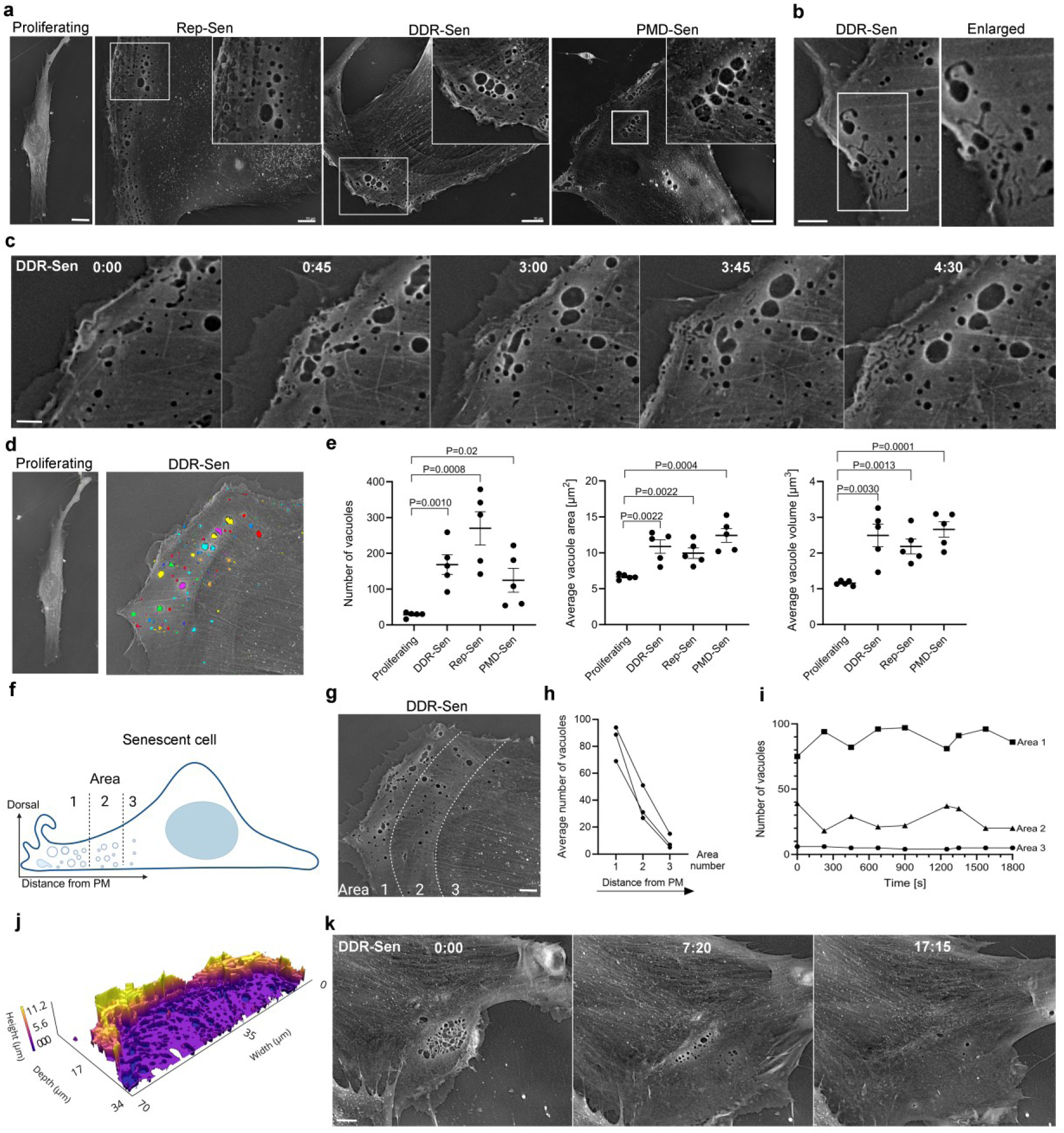
Spatiotemporal dynamics of vacuole formation in senescent cells. **(a)** Representative live cell images captured by holotomography (Tomocube) show cytoplasmic vacuoles (white arrows) and membrane ruffling in proliferating and senescent WI-38 cells. Scale bar 10 μm. **(b)** Representative live cell image of DDR-Sen WI-38 cell showing tubular shape of newly formed vacuoles. Scale bar 10 μm. **(c)** Representative ROI from holotomography timelapse video recording (Movie 5) of vacuole maturation process in DDR-Sen WI-38 cell. Scale bar 5 μm. **(d)** Example of image preparation in TomoAnalysis software for vacuole quantification **(e)** Quantification of vacuole number, average area, and average volume in proliferating and senescent WI-38 cells. n=5. **(f)** Example of image segmentation to Area 1-3. **(g)** Scheme illustrating the ROI-analysis used to quantify vacuoles with increasing distance from the plasma membrane. Scale bar 10 μm. **(h)** Quantification of vacuole number in DDR-Sen cells in each area. n=3 **(i)** Quantification of vacuole number in DDR-Sen cell in each area with time. n=1. **(j)** Reconstructed 3D heatmap from refractive index showing wave-like dorsal ruffles in senescent cell. **(k)** Representative frames from timelapse video recording (Movie 8) of vacuole disappearance in DDR-Sen WI-38 cell. Scale bar 10 μm.

To assess the spatial distribution of vacuoles, we subdivided the peripheral cytoplasmic region of DDR-Sen cells into three concentric zones (Areas 1–3) based on increasing distance from the plasma membrane, extending up to 45 µm (Fig. 2f and g). We show that although the total number of vesicles decreased with increasing distance from the plasma membrane, the number of vacuoles within each region remained relatively stable over time (Fig. 2g-i). This observation raises the possibility that vacuoles move in a coordinated manner along cytoskeletal structures, rather than separate stochastic trafficking. Notably, vacuoles emerged after dorsal, wave-like membrane structures, with height reaching 11 µm (Fig. 2j) and were no longer detectable after reaching the curved actin bundles positioned perpendicular to the lamellar network (Fig. 2k and Movie 8). This observation suggests that interactions with specific actin architecture may regulate the retention, movement, or disassembly of these vacuoles. To further validate that these structures might represent macropinosomes, we performed live-cell imaging of stable cells expressing Phafin2-GFP or the GFP-2xFYVE domain, two established markers of nascent and maturing macropinosomes (Movies 9 and 10). Both markers revealed the emergence of large vesicles at the plasma membrane followed by their maturation, centripetal trafficking, and eventual disappearance, recapitulating the dynamics observed by holotomographic imaging.

Together, these observations indicate that in senescent cells, vacuoles form at the cell periphery and undergo inward movement accompanied by morphological and size changes, ultimately disappearing near transverse arc-like actin structures.

### Macropinocytosis is upregulated in senescent cells

Cytoplasmic vacuoles observed in senescent WI-38, BJ, and U2OS cells showed morphological hallmarks of macropinosomes, including their large size (>2 μm²), spherical appearance, characteristic peripheral spatial arrangement and Rab8, 21, 5, FYVE-domain and Phafin2 identity^62,63^. To determine whether these structures are *bona fide* macropinosomes, we incubated the cells with high-molecular weight dextran conjugated to either FITC or Texas Red, a well-established macropinocytosis cargo^45,46,48^. Both confocal microscopy and 3D holotomographic imaging revealed robust dextran accumulation within large intracellular vesicles of senescent cells (Supplementary Fig. 2a and b). Co-localization of dextran with internalized WGA further confirmed the macropinosomal identity of these vesicles (Supplementary Fig. 2b). Next, we compared the dextran internalization capacity of senescent and proliferating cells. WI-38, BJ, and U2OS cells were incubated with dextran for 60 minutes. This showed significantly higher dextran uptake in senescent cells, whereas proliferating cells exhibited minimal incorporation (Fig. 3a and b, Supplementary Fig. 2c). Importantly, dextran uptake increased progressively with the senescence phenotype. To validate the establishment of senescence over time, we induced senescence in WI-38 cells and monitored the progressive decrease of Lamin B1 expression by western blotting (Supplementary Fig. 2d and e). In parallel, cells were incubated with dextran at days 4, 8, and 16 following senescence induction. Quantification of dextran uptake at these time points revealed a time-dependent upregulation of macropinocytosis (Supplementary Fig. 2f and g). These findings suggest that macropinocytosis increases during the progression of senescence.

**Figure 3.**
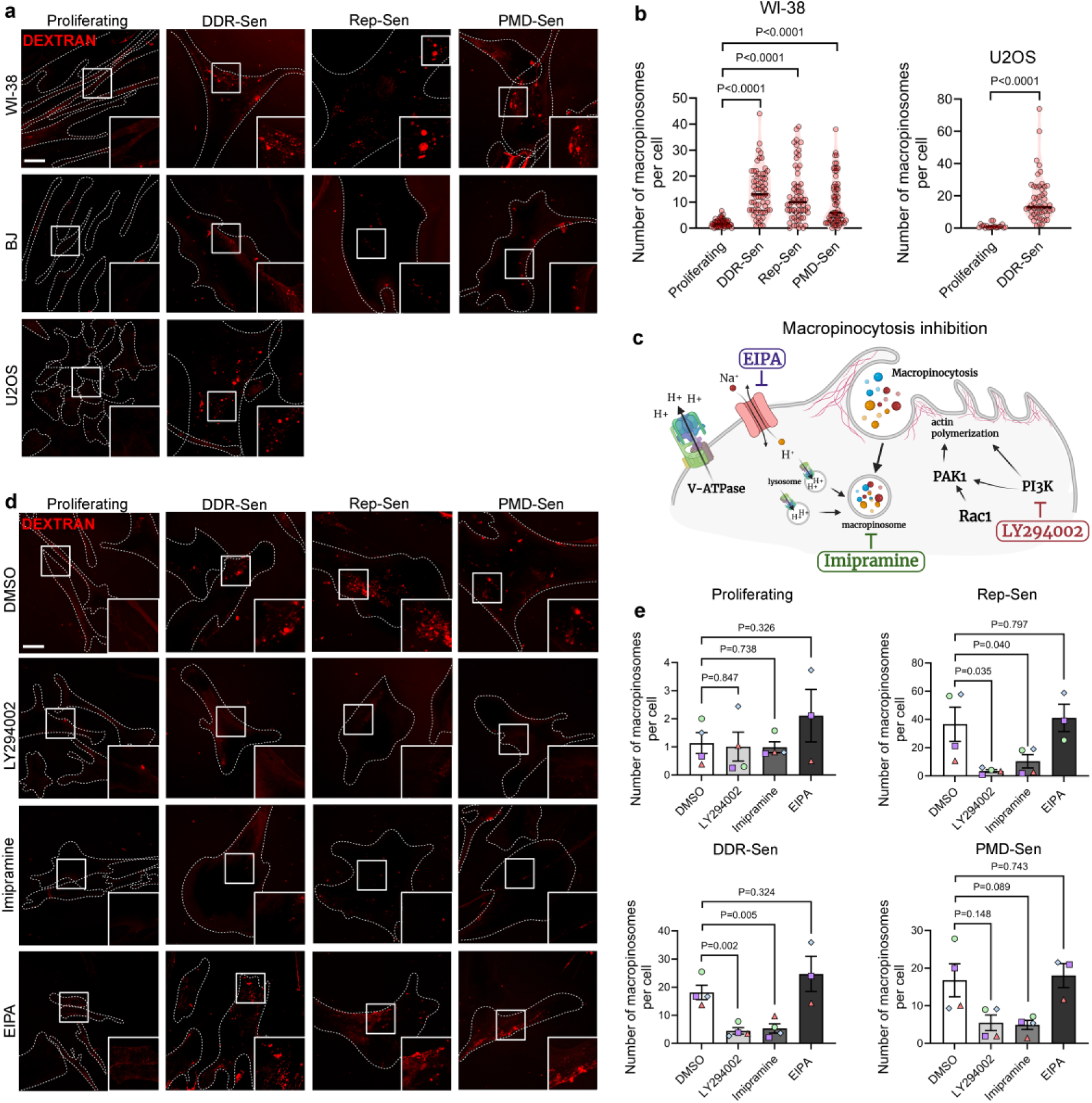
Senescent cells dynamically form new large vesicles through macropinocytosis. **(a)** Macropinocytosis assay using Texas Red-dextran (red) indicates that senescent WI-38 cells display increased levels of macropinocytosis compared to proliferating cells. Scale bar 20 μm. **(b)** Quantification of macropinocytic uptake in WI38 and U2OS cells. **(c)** Graphical illustration of simplified macropinocytosis mechanism and inhibition strategy. **(d)** Confocal microscopy analysis of fluorescent dextran uptake in proliferating and senescent WI-38 cells. Cells were pretreated with DMSO, LY294002, Imipramine or EIPA for 30 min. **(e)** Quantitative analysis of images in d. n=4.

Macropinocytosis is typically amiloride sensitive and phosphatidylinositol 3-kinase (PI3K)-dependent^46^. To further assess whether the vacuoles in senescent cells meet these criteria, we examined the sensitivity of dextran uptake to the macropinocytosis pharmacological inhibitors^45^ (Fig. 3c). Senescent WI-38 cells were pre-treated with each inhibitor for 30 min, followed by dextran loading for 60 min. Inhibitors were present in the medium during an entire experiment. Treatment with LY294002 (PI3K inhibitor) and imipramine (macropinocytosis inhibitor) reduced dextran internalization across all senescent subtypes (Fig. 3c-e). In contrast, EIPA, an inhibitor of Na⁺/H⁺ exchangers widely used to block macropinocytosis, did not reduce dextran uptake in senescent cells (Fig. 3c-e). This EIPA resistance aligns with other reports in immune cells, such as immature dendritic cells and macrophages, in which amiloride derivatives similarly fail to inhibit macropinocytosis^45^.

Collectively, these results suggest that macropinocytosis is upregulated in senescent cells and shows sensitivity to PI3K and actin polymerization inhibition and resistance to the pH-regulatory mechanisms associated with amiloride-sensitive macropinocytosis.

### Macropinocytosis in senescent cells is glutamine and Ca^2+^ sensitive but EGF-independent

Macropinocytosis is a growth factor-induced endocytic process, although immune cells, such as immature dendritic cells and macrophages, also exhibit a constitutive, stimulus-independent macropinocytosis^45,65^. To determine whether macropinocytosis in senescent cells is constitutive or relies on external stimulation, we first assessed dextran uptake in Ca^2+^-depleted medium. We show that short Ca^2+^ depletion (30 min pre-treatment in Ca²⁺-free DMEM) reduced dextran uptake in senescent cells, suggesting that Ca²⁺ contributes to efficient macropinocytosis (Fig. 4a and b, Supplementary Fig. 3a). Next, we assessed serum-starved conditions. Senescent WI-38 cells were incubated in medium without FBS for 16 hours. We show that senescent cells internalized dextran even in the absence of serum, whereas internalized dextran was undetectable in proliferating controls (Fig. 4c and d, No FBS). These findings suggest that senescent cells, unlike their proliferating counterparts, show serum-independent, constitutive macropinocytosis. Consistently, stimulation with epidermal growth factor (EGF) (100 ng/mL for 60 min) did not further change dextran uptake in senescent cells when compared to medium without serum (Fig. 4c and d, No FBS vs No FBS, EGF), despite activation of EGF endocytosis in proliferating cells (Supplementary Fig. 3b) and canonical downstream EGF signaling verified by western blotting (Supplementary Fig. 3c). This implies that macropinocytosis in senescent cells does not respond to additional growth factor stimulation, showing features of constitutive, growth factor-independent uptake. We next examined how other environmental cues regulate macropinocytosis in senescent cells. Glutamine withdrawal is known to induce macropinocytosis particularly in cancer cells where it supports cell survival^46^. Unlike Ca^2+^ depletion or EGF stimulation, glutamine starvation (-Q, 24 h in glutamine-free DMEM) further increased dextran internalization specifically in Rep-Sen WI-38 cells (Fig. 4e and f). Notably, this condition also elevated p-AMPKα levels only in senescent WI-38 cells, but not in proliferating counterparts (Fig. 4g and Supplementary Fig. 3d).

**Figure 4.**
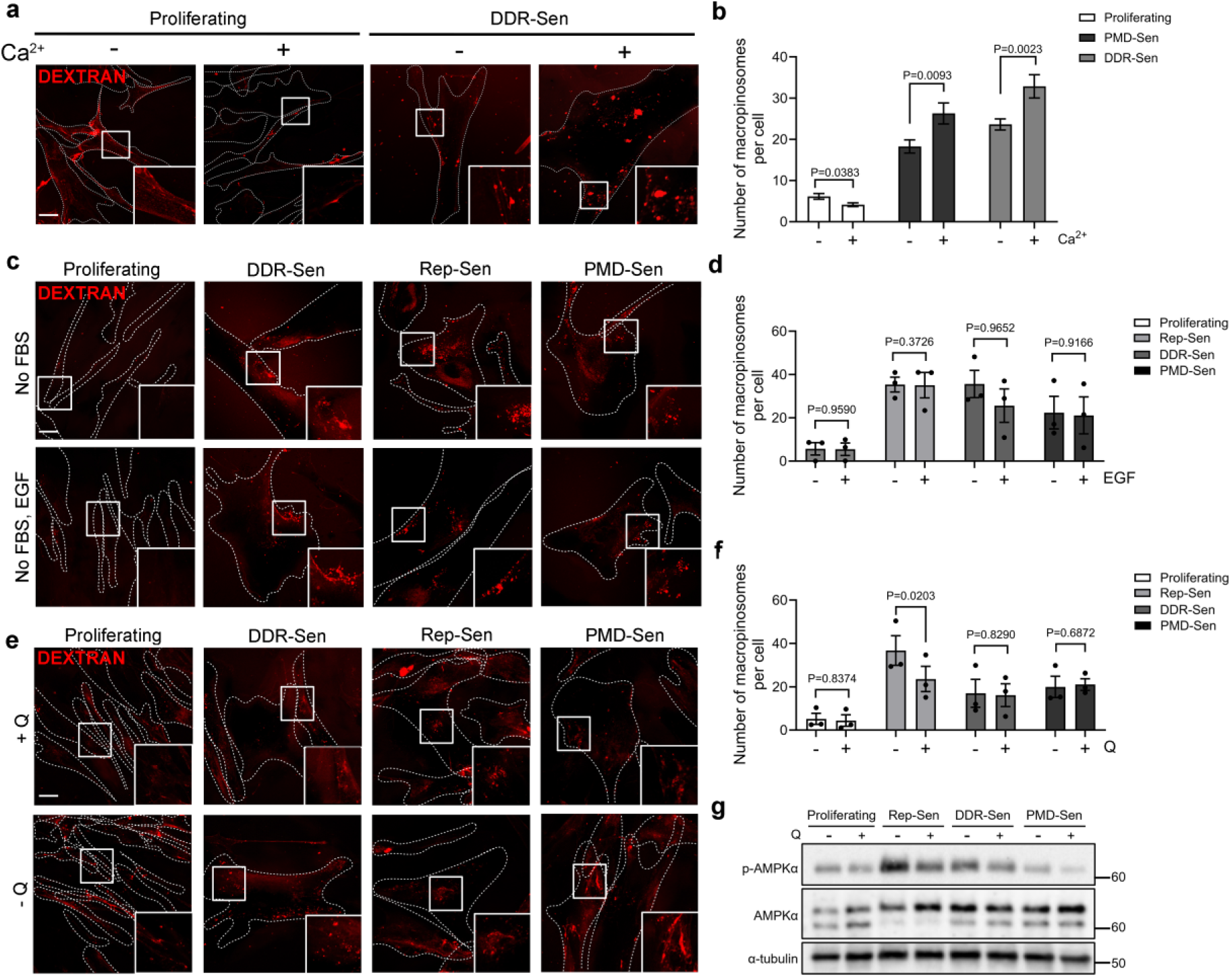
Macropinocytosis in senescent cells is calcium and glutamine sensitive and EGF-independent. **(a)** Confocal microscopy analysis of fluorescent dextran uptake in proliferating and senescent WI-38 cells. Cells were incubated in medium without extracellular calcium 30 min before dextran loading. Scale bar 20 μm**. (b**) Quantification of macropinocytic uptake in a. n=3 **(c)** Confocal microscopy analysis of fluorescent dextran uptake in proliferating and senescent WI-38 cells. Cells were incubated in medium without FBS for 16h. Were indicated, EGF was added for 60 min. Scale bar 20 μm. **(d)** Quantification of macropinocytic uptake in c. n=3. **(e)** Confocal microscopy analysis of fluorescent dextran uptake in proliferating and senescent WI-38 cells. Cells were incubated in medium without glutamine (-Q) for 24h. Scale bar 20 μm. **(f)** Quantification of macropinocytic uptake in e. n=3. **(g)** Western blotting using lysates of proliferating and senescent WI-38 cells. The lysates were collected 16 days after treatment with SDS or doxorubicin and after 24h incubation in media with or without Q.

Taken together, these findings suggest that macropinocytosis in senescent cells is growth factor-independent and resembles the constitutive fluid-phase internalization in dendritic immune cells^45^. However, in glutamine-deprived conditions, macropinocytosis in senescent cells parallels the adaptive response observed in cancer cells and CAFs, that is regulated through AMPKα signaling^48^.

### PAK1 regulates macropinocytosis in senescent cells

p21-activated kinase 1 (PAK1) is a well-established effector downstream of PI3K and Rac1-Cdc42 signaling, and it regulates actin cytoskeletal remodeling, membrane ruffling, and macropinosome closure^66^. Given its canonical role in macropinocytic cup formation, scission and trafficking, we hypothesized that PAK1 contributes to the upregulated macropinocytic activity of senescent cells. To examine the spatial distribution of PAK1, we transduced U2OS cells with a lentiviral construct encoding GFP-tagged PAK1 and assessed its localization by fluorescence microscopy. Senescent cells exhibited pronounced GFP-PAK1 enrichment at the cell periphery, compared to their proliferating counterparts (Fig. 5a). This localization was consistent with our hypothesis that PAK1 may regulate macropinocytic membrane remodeling in senescent cells. To further test this possibility, we next pharmacologically inhibited PAK1 using IPA-3, a selective allosteric inhibitor that prevents PAK1 activation by blocking its interaction with upstream GTPases^67^.

**Figure 5.**
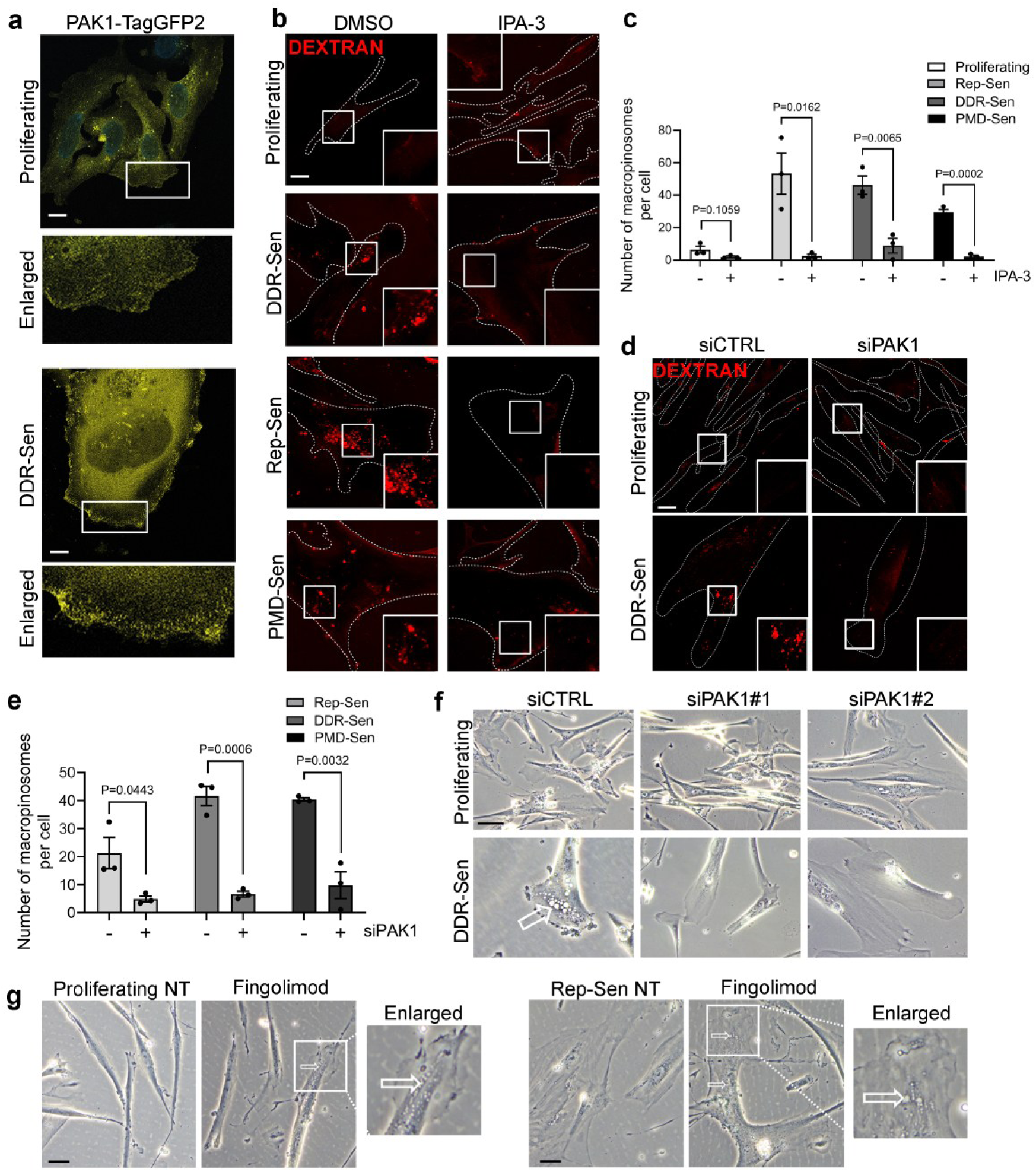
PAK1 is required for macropinocytosis in senescent cells. **(a)** Localization of PAK1 in U2OS proliferating and senescent cells expressing PAK1-TagGFP2. **(b)** Confocal microscopy analysis of fluorescent dextran uptake in proliferating and senescent WI-38 cells. Cells were pretreated with IPA-3 30 min before dextran loading. Scale bar 20 μm. **(c)** Quantification of macropinocytic uptake in b. n=3. **(d)** Confocal microscopy analysis of fluorescent dextran uptake in proliferating and senescent WI-38 cells. Cells loaded with dextran for 45min after PAK1 knockdown for 72h. Scale bar 20 μm. **(e)** Quantification of macropinocytic uptake in d. n=3. **(f)** Microscopy analysis of vacuole formation in proliferating and senescent WI-38 cells with and without PAK1 knockdown. **(g)** Microscopy analysis of vacuole formation in proliferating and senescent WI-38 cells after stimulation with fingolimod for 3H.

Proliferating and senescent WI-38 cells were pre-treated with IPA-3 for 30 min, followed by a 45-min dextran uptake assay. IPA-3 reduced dextran internalization in senescent WI-38 cells indicating that PAK1 activity is required for macropinocytosis in senescent cells (Fig. 5b and c). We next depleted PAK1 using siRNA-mediated knockdown (Supplementary Fig. 4a). PAK1 knockdown not only significantly downregulated dextran uptake but also reduced the number of senescent cells forming cytoplasmic vacuoles (Fig. 5d-f and Supplementary Fig. 4b). PAK1 knockdown similarly reduced dextran uptake in senescent U2OS cells (Supplementary Fig. 4c). These combined results show that PAK1 is necessary for macropinosome formation and accumulation in senescent cells.

Next, we asked whether PAK1 activation alone is sufficient to induce macropinocytosis. To this end, we utilized fingolimod (FTY720), a sphingosine-1-phosphate (S1P) receptor modulator known to stimulate PAK1 indirectly via Rac1^68,69^. Short-term fingolimod treatment (5 μM, 3 h) induced pronounced vacuole formation in both proliferating and senescent WI-38 cells (Fig. 5g). This cellular response indicated that PAK1 activation is sufficient to drive the formation of macropinosomes and promote fluid-phase uptake.

These findings identify PAK1 as a regulator of macropinocytosis and vacuole formation in senescent cells. Both loss-of-function and gain-of-function approaches converge to show that PAK1 regulates the vesicle formation machinery that drives senescence-associated macropinocytosis.

### PAK1 is dispensable for senescence induction and maintenance

To determine if PAK1 is required for the induction or maintenance of cellular senescence, we systematically depleted PAK1 at multiple stages of the senescence progression (Fig. 6a). Knockdown was performed prior to senescence induction, at early timepoints following DNA damage or plasma membrane damage (day 4), at a late stage (day 16), or in already established replicative senescence. This design enabled us to assess the role of PAK1 in the initiation of senescence and maintaining the senescent phenotype.

**Figure 6.**
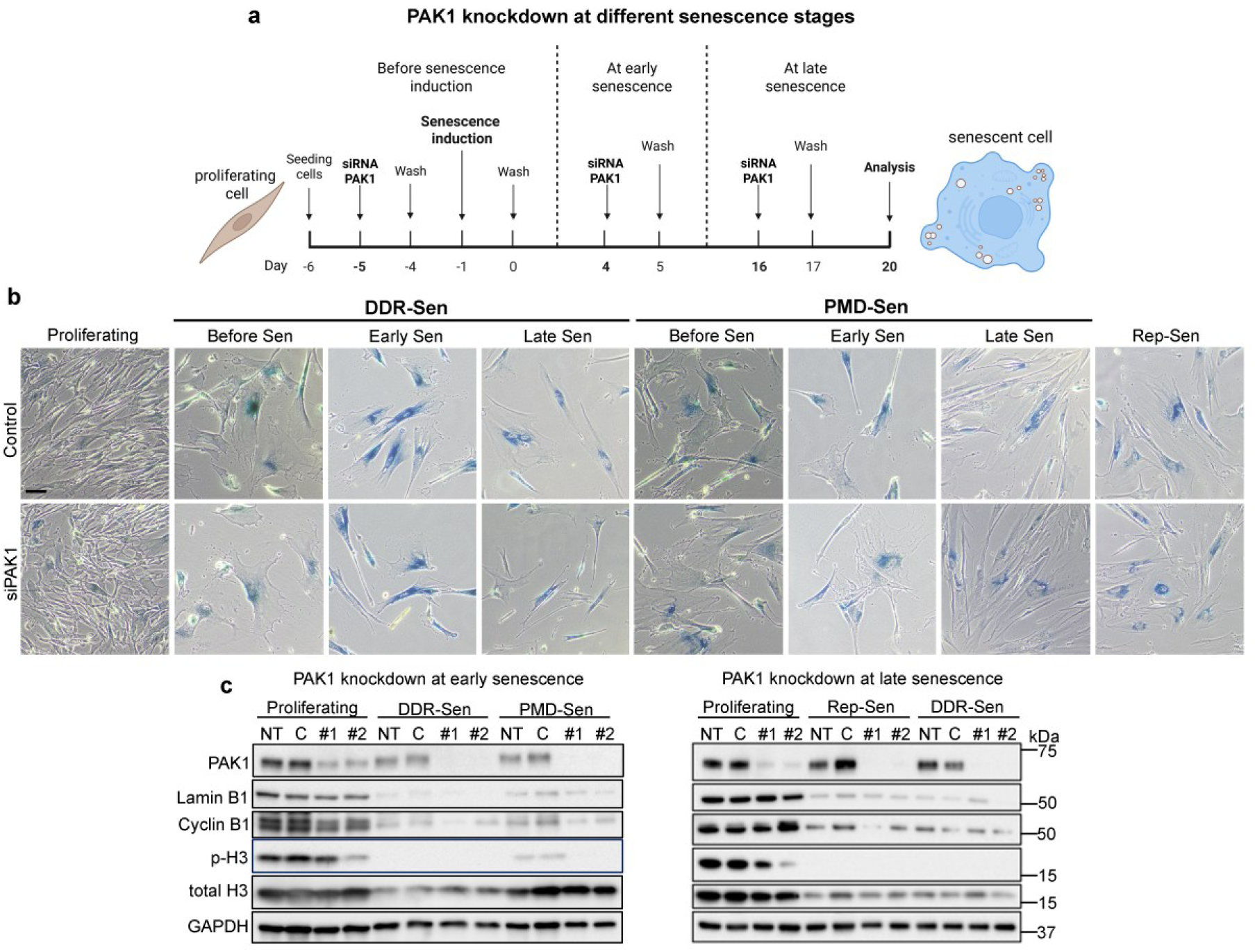
PAK1 knockdown does not alter senescence induction and maintenance in WI-38 cells. **(a)** Scheme of PAK1 depletion strategy at different senescence stages. **(b)** SA-β-Gal staining of WI-38 with and without PAK1 knockdown at different senescence day. **(c)** Expression of PAK1, senescence marker Lamin B1 and proliferation markers cyclin B1 and histone H3 in proliferating and senescent cells upon PAK1 depletion by western blotting. NT – not treated, C – siScramble control, #1 – siPAK1#1, #2 – siPAK1#2.

Across all senescence subtypes tested, canonical markers of senescence remained largely unaffected by PAK1 depletion (Fig. 6b and c). SA-β-Gal staining showed no detectable difference between PAK1-depleted and control cells in all conditions tested here (Fig. 6b). These results suggest that knockdown of PAK1, whether early or late, does not attenuate the SA-β-Gal-positive phenotype. Consistently, Lamin B1 decrease was comparable between PAK1-knockdown and control senescent cells (Fig. 6c) These results suggest that the establishment and maintenance of cellular senescence are independent of PAK1. We next examined cell-cycle exit. Following induction of senescence, both cyclin B1 and phosphorylated histone H3 (p-H3), the markers of G2/M progression and mitotic entry, respectively, were downregulated with or without PAK1 knockdown (Fig. 6c). The decline in these mitotic markers across all conditions tested here suggests that PAK1 is dispensable for the establishment and maintenance of cell-cycle arrest.

### Knockdown of PAK1 suppresses inflammatory SASP, IL6, CCL2, and IL12A, via regulation of TGFβ/SMAD signaling in senescent cells

To investigate the transcriptional changes induced by PAK1 knockdown, we performed RNA sequencing on proliferating and Rep-Sen WI-38 cells with and without siRNA-mediated PAK1 knockdown. Principal component analysis (PCA) and distance matrices showed that PAK1 depletion induced transcriptomic alterations in both proliferating and senescent cells; however, the magnitude of transcriptional reprogramming was higher in the replicative senescent cells than in proliferating cells (Fig. 7a and Supplementary Fig. 5a). Differential expression profiles further suggested that the consequence of PAK1 knockdown was context-dependent, i.e., proliferating and senescent cells showed distinct sets of altered genes and pathways (Supplementary Fig. 5b and c). As a control for knockdown efficiency and senescence induction in the RNA-seq samples, we examined gene expression levels (logCPM) of PAK1 and p16 (CDKN2A) across the groups. PAK1 expression was reduced in the PAK1 knockdown groups, while p16 expression was elevated in replicative senescent (Rep-Sen) samples compared to proliferating cells (Supplementary Fig. 5d).

**Figure 7.**
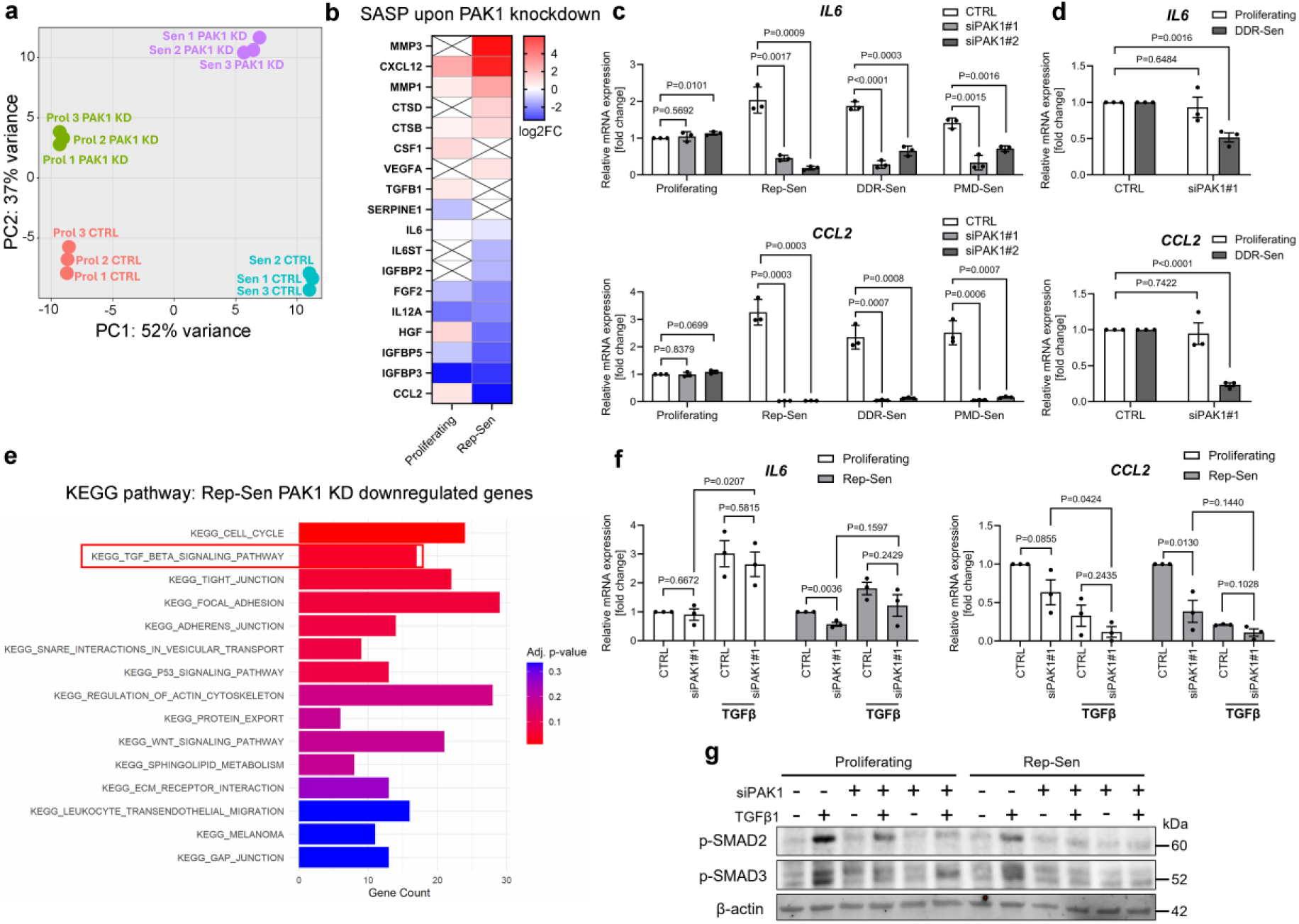
PAK1 depletion suppresses proinflammatory SASP gene expression in senescent cells through TGFβ signaling. **(a)** PCA of RNA-seq samples showing separation of control and PAK1-depleted cells in proliferating (Prol) and Rep-Sen (Sen) WI-38 cells. **(b)** Heatmap showing RNA-seq–derived changes in SASP-related gene expression in proliferating and Rep-Sen WI-38 cells upon PAK1 depletion. **(c, d)** qPCR analysis of *IL6* and *CCL2* expression in proliferating and senescent WI-38 (c) and U2OS (d) cells following PAK1 knockdown with two different siRNA (siPAK1#1 and siPAK1#2). **(e)** KEGG pathway analysis of only significantly downregulated DEGs (log2FC < 1, p-value < 0.05) in the Rep-Sen PAK1 knockdown group. **(f)** qPCR analysis of *IL6* and *CCL2* expression following treatment with recombinant TGFβ1 (5 ng/mL, 24 h) in control and PAK1-depleted proliferating and Rep-Sen WI-38 cells. **(g)** Immunoblot analysis of phosphorylated SMAD2 and SMAD3 in proliferating and senescent cells following PAK1 depletion and/or TGFβ1 treatment, demonstrating reduced SMAD2/3 activation in PAK1-depleted cells. Two different siPAK1 sequences were used to knockdown PAK1.

A consequence of PAK1 knockdown in senescent cells was the suppression of the canonical proinflammatory SASP factors (Fig. 7b). SASP cytokines, including *IL6*, *CCL2*, and *IL12A*, were significantly downregulated at the transcript level, a finding validated by qPCR across multiple models of senescence in WI-38 (Fig. 7c and Supplementary Fig. 5e) and in U2OS cells (Fig. 7d). Notably, pharmacological inhibition of PAK1 using IPA-3 and NVS-PAK1-1 recapitulated these effects, yet to a lesser extent, confirming that PAK1 kinase activity contributes to SASP maintenance (Supplementary Fig. 5f).

KEGG pathway analyses identified TGFβ signaling as one of the most significantly downregulated pathways following PAK1 knockdown in senescent but not proliferating cells (Fig. 7e and Supplementary Fig. 5g). Examination of individual TGFβ pathway components revealed divergent responses in proliferating and senescent cells (Supplementary Fig. 5h). In proliferating cells, PAK1 depletion modestly changed TGFβ pathway activity, with the upregulation of expression of *TGFB1*, *TGFBR3*, *ACVR1B*, *ACVR2B*, and the extracellular matrix gene *COL3A1*. In contrast, senescent PAK1-depleted cells displayed attenuation of TGFβ/SMAD signaling, including reduced expression of *TGFB2*, *TGFBR2*, *TGFBR3*, *SMAD2*, and *SMAD3*, together with upregulation of the inhibitory feedback mediator *SMAD7*. These results suggest the presence of negative feedback that selectively downregulates TGFβ signaling in senescent cells. qPCR analysis results were consistent with the RNA-seq results, confirming downregulation of *TGFB2* and *SMAD3* and upregulation of *SMAD7* following PAK1 depletion in senescent WI-38 cells (Supplementary Fig. 5i).

To assess whether suppression of TGFβ signaling contributes to the suppressed SASP, we performed rescue experiments using exogenous TGFβ1 (5 ng/mL for 24 h) under PAK1 depletion condition. TGFβ1 treatment alone enhanced *IL6* and decreased *CCL2* expression in control senescent cells, showing different effects of TGFβ in SASP regulation (Fig. 7f). Notably, in PAK1-depleted cells, TGFβ1 partially restored *IL6* expression while further suppressing *CCL2*, indicating that at least part of the PAK1-dependent SASP phenotype is mediated through TGFβ/SMAD signaling (Fig. 7f). The immunoblot analysis showed reduced phosphorylation of SMAD2 and SMAD3 in PAK1-depleted senescent cells, with only partial downregulation observed in proliferating cells (Fig. 7g). These results confirm that PAK1 knockdown attenuates TGFβ pathway at the mRNA and protein levels in senescent cells. The transcriptomic profiling, targeted validation, chemical inhibition, and rescue experiments converge on a model that PAK1 supports the inflammatory phenotype of senescent cells partially by maintaining TGFβ/SMAD signaling. PAK1 depletion suppressed *IL6, CCL2, IL12A* gene expression and TGFβ pathway activity in senescent cells.

### Macropinocytosis contributes to SASP regulation

To determine whether macropinocytosis contributes to SASP regulation, or if PAK1 regulates SASP independently of macropinocytosis, we assessed the contribution of macropinocytosis structural machinery to SASP. To this end, we disrupted cytoskeletal dynamics using latrunculin A (LatA) and cytochalasin D (CytoD). Both compounds interfere with the actin polymerization necessary for membrane ruffling, cup formation, and macropinosome closure. Pre-treatment of WI-38 cells with either inhibitor resulted in significantly inhibited dextran uptake (Fig. 8a and b) and downregulated expression of *IL6* (Fig. 8c and Supplementary Fig. 6a) in senescent but not in proliferating cells. Expression of *CCL2* decreased after treatment with LatA but not with CytoD in senescent cells under the conditions tested (Fig. 8c and Supplementary Fig. 6a). Moreover, to determine whether this effect extends beyond disruption of actin dynamics, we targeted additional PAK1-independent components of the macropinocytosis machinery. We targeted Phafin2, a key regulator of macropinosome maturation, and depleted it using siRNA, and we used imipramine, an inhibitor of acid sphingomyelinase that impairs macropinocytosis. Both interventions significantly reduced *IL6* and *CCL2* expression in senescent U2OS cells (Fig. 8 c and d). Since senescent U2OS cells exhibit increased expression of not only *IL6* and *CCL2* but also other SASPs, namely *IL8* and *IL1β* (Supplementary Fig. 6b), we next examined these cytokines under macropinocytosis inhibition conditions. Both *IL8* and *IL1β* were robustly downregulated following Phafin2 depletion and imipramine treatment in senescent cells (Supplementary Fig. 6c-e).

**Figure 8.**
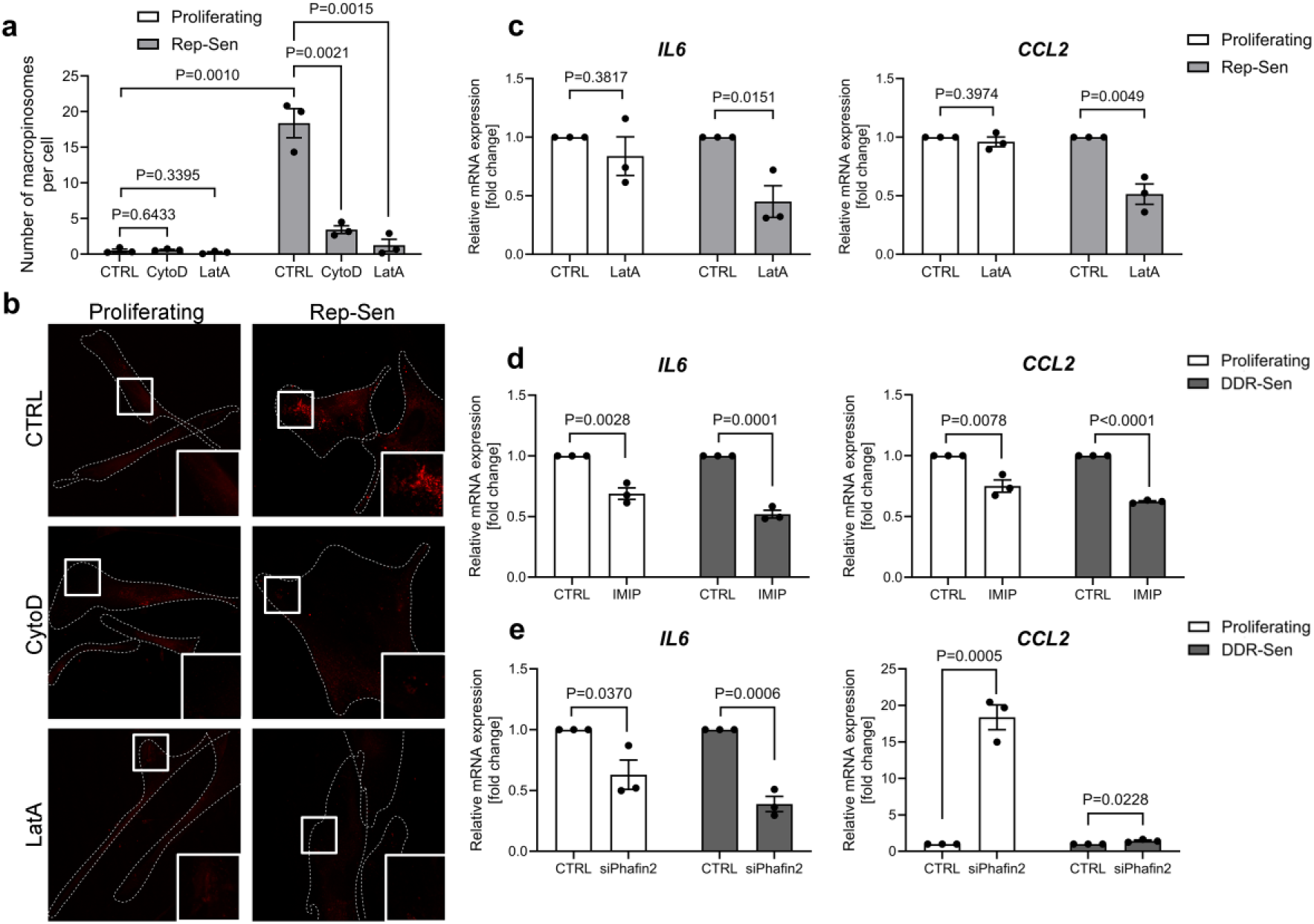
Macropinocytosis contributes to SASP expression. **(a)** Quantification of dextran uptake in proliferating and Rep-sen WI-38 cells after pretreatment (1h) with actin polymerization inhibitors, Cytochalasin D (CytoD) or Latranculin A (LatA). n=3 **(b)** Confocal microscopy analysis of fluorescent dextran uptake in proliferating and senescent WI-38 cells. Scale bar 20 μm. **(c)** qPCR analysis of *IL6* or *CCL2* expression following treatment with LatA in proliferating and Rep-Sen WI-38 cells. n=3. **(d)** qPCR analysis of *IL6* or *CCL2* expression following treatment with imipramine in proliferating and DDR-Sen U2OS cells. n=3 **(e)** qPCR analysis of *IL6* or *CCL2* expression following Phafin2 siRNA-mediated knock down in proliferating and DDR-Sen U2OS cells. n=3

Finally, we investigated whether inhibition of Rac1, upstream activator of PAK1 and an essential regulator of membrane ruffling during macropinocytosis, similarly affects SASP expression. Rac1 knockdown significantly reduced expression of *IL6*, *CCL2*, *IL8*, and *IL1β* in senescent U2OS cells (Supplementary Fig. 6 d and f). Together, these findings confirm that inhibition of multiple distinct components of macropinocytosis leads to suppression of inflammatory SASP gene expression in senescent cells.

## Discussion

Here we report that replicative and stress-induced senescent WI-38 and BJ fibroblasts, and U2OS osteosarcoma cells show pronounced peripheral cytoplasmic vacuole formation. Using brightfield, confocal, and holotomography microscopy, we demonstrated that these vacuoles are formed through upregulated macropinocytosis. Based on dextran uptake assays, we found that macropinocytosis in senescent cells is PI3K-dependent, regulated by Ca^2+^ and glutamine availability, and occurs independently of growth factor stimulation. We further identified PAK1 as a key regulator of macropinocytosis in senescent cells, as it controlled dextran uptake and macropinosome formation. However, PAK1 knockdown did not affect the establishment or maintenance of senescence. Finally, we showed that macropinocytosis contributes to the proinflammatory SASP expression, including *IL6*, *CCL2, IL8* and *IL1β*. Together, our work provides mechanistic insight into macropinocytosis in senescence and uncovers its role in regulating SASP.

Macropinocytosis is an actin-dependent endocytic pathway characterized by membrane ruffling and formation of large vesicles termed macropinosomes, which enable non-selective bulk uptake of extracellular fluid together with nutrients, plasma membrane, receptors and signaling molecules^70,71^. In the present study, we found that macropinocytosis progressively increases following senescence induction, as reflected by rising dextran uptake as cells transition toward a fully senescent state (Supplementary Fig. 2d-g). This gradual increase aligns with the growing recognition that senescence is not a binary state but a dynamic and evolving phenotype that continues to develop after induction, with progressive transcriptional, proteomic, secretory (SASP), and morphological changes^17,72^. Our findings suggest that enhanced macropinocytosis also arises as a consequence of the developing senescence phenotype.

A widely studied form of macropinocytosis is growth factor-induced macropinocytosis, observed in fibroblasts, epithelial cells and many types of cancers^46,48^. In contrast, Canton et al. reported constitutive macropinocytosis performed in antigen-presenting immune cells that occurs without additional stimulation^45^. Consistent with these findings, we show here that senescent cells internalize dextran constitutively in the absence of growth factors and do not respond to EGF stimulation (Fig. 4c and d), indicating an epidermal growth factor-independent uptake or growth factor hyposensitivity. In line with this observation, we further show that macropinocytosis in senescent cells is regulated by ionic cues, as calcium ion (Ca^2+^) depletion impaired dextran uptake (Fig. 4a and b). This sensitivity to Ca^2+^ in the medium resembles CaSR-dependent macropinocytosis observed in immune cells and suggests that constitutive macropinocytosis may represent a conserved regulatory mechanism in non-proliferative or specialized cellular states^45^. Notably, replicative senescent cells remain responsive to glutamine (Q) depletion, further enhancing macropinocytosis under -Q conditions and activating AMPKα signaling (Fig. 4e-g). A similar dependency on macropinocytosis for Q acquisition and survival has been described in cancer cells and cancer-associated fibroblasts under nutrient-limited conditions^48^. Our findings therefore suggest that macropinocytosis in senescent cells is constitutive while remaining selectively responsive to changes in extracellular nutrient availability.

Building upon prior identification of vacuoles in senescent cells^41,42,57–59^ our study advances mechanistic understanding by outlining the role of PAK1 as a non-redundant regulator of macropinocytosis in senescent cells. While recent work by Nagano et al.^41,42^ established macropinocytosis in senescence and linked it to LY6D and integrin-mediated signaling pathways, our results directly implicate PAK1, kinase important for actin polymerization, orchestrating membrane ruffle and vacuole formation. Pharmacological inhibition of PAK1 using IPA-3 and NVS-PAK1-1 and siRNA-mediated knockdown reduced macropinocytic uptake and vacuole formation, underscoring its necessity (Fig. 5b-f and Supplementary Fig. a-c). Conversely, activation of PAK1 with fingolimod induced vacuole generation even in proliferating cells, indicating sufficiency (Fig. 5g). These findings position PAK1 as a master regulator integrating upstream signals to the cytoskeletal and membrane remodeling machinery required for macropinocytosis in senescent cells.

In this study, we performed transcriptomic analyses of PAK1 knockdown in WI-38 replicative senescent and proliferating cells. We revealed that PAK1 knockdown leads to a downregulation of proinflammatory SASP factors in senescent cells, including *IL6*, *CCL2* and *IL12A*, alongside decreased expression of TGFβ receptors (*TGFBR2* and *TGFBR3*) and *SMAD2/3*, the intracellular effectors of TGFβ signaling (Fig. 7b-d and Supplementary Fig. 5e). Conversely, negative feedback regulator of TGFβ signaling *SMAD7* was upregulated, suggesting that PAK1 maintains a positive feed-forward loop. TGFβ signaling can act as a tumor suppressor by enforcing growth arrest, yet it also promotes chronic proinflammatory environment depending on cellular context and senescence subtype^73,74^. In addition, several studies have shown that TGFβ signaling contributes to paracrine senescence, whereby TGFβ secreted by primary senescent cells induces growth arrest and senescence-associated phenotypes in neighboring cells, amplifying tissue-level responses^75,76^. Given that endocytosis can modulate receptor availability and downstream signaling, through receptor internalization and recycling, it is plausible that PAK1-mediated macropinocytosis influences TGFβ receptor trafficking or localization, sustaining pathway activity. Future studies dissecting how PAK1 interfaces with TGFβ will be critical to fully understand this regulatory axis.

The role of PAK1 in cellular aging and longevity has been investigated in model organisms, notably *Caenorhabditis elegans*. Yanase et al. ^56^ reported that loss of PAK1 extended lifespan in *C. elegans*, suggesting a role for PAK1 in aging regulation. In this context, PAK1 functions by suppressing the activity of DAF-16, the worm ortholog of the mammalian FOXO transcription factors, which are key regulators of stress resistance and longevity. Moreover, in our mammalian cell system, PAK1 knockdown did not alter classical hallmarks of senescence such as expression of mitotic markers and SA-β-Gal, suggesting that PAK1 is not required for mitotic exit or maintenance of growth arrest during senescence (Fig 6). Instead, our findings showed that PAK1 plays a role in modulating SASP. This suggests that PAK1 may influence aging not by controlling cell cycle arrest, but rather by modulating metabolic and inflammatory outputs.

In this study we show that macropinocytosis contributes to the inflammatory phenotype of senescent cells. Disruption of macropinocytosis through PAK1-independent actin polymerization suppression, decreased *IL6* and *CCL2* expression. In addition to reducing *IL6* and *CCL2*, inhibition of the macropinosome maturation factor Phafin2, treatment with imipramine, or depletion of Rac1 also decreased *IL8* and *IL1β* expression, showing that targeting multiple components of the macropinocytosis pathway consistently suppresses inflammatory SASP gene expression in senescent cells. One possible explanation is that macropinocytosis supports internalization and trafficking of signaling components necessary for SASP gene expression, through receptor and plasma membrane turnover. Consistent with this idea, increased macropinocytosis has recently been reported^77^ in mouse fibrotic system, where macropinocytosis inhibition reduced TGFβ-regulated profibrotic transcriptional programs. Given that senescent cells accumulate in fibrotic tissues and contribute to chronic inflammatory signaling, these observations suggest a potential mechanistic intersection between senescence-associated macropinocytosis and fibrotic diseases.

In conclusion, our results highlight upregulated macropinocytosis in senescent cells that is constitutive, glutamine- and Ca^2+^-sensitive, and regulated by PAK1. Our findings redefine macropinocytosis as a regulator of the proinflammatory SASP and position PAK1 as a promising therapeutic target for modulating senescence-caused chronic inflammation and tissue deterioration. Future studies should dissect how vesicular trafficking networks intersect with canonical signaling pathways to orchestrate the complex SASP landscape. Moreover, exploring whether senescent subtypes with differential macropinocytic capacities exist in vivo, and how they impact tissue aging and organismal health, will be critical.

## Methods

### Cell lines and cell culture

WI-38, BJ and U2OS cells (WI-38 cells, fibroblast, female, PDL 36.6 from RIKEN BRC, cat. no. RCB0702, RRID: CVCL_0579; WI-38 cells, fibroblast, female, PDL 32.2 from JCRB, cat. no. IFO50075, RRID: CVCL_0579; BJ cells, fibroblast, male, PDL 24.0 from the American Type Culture Collection, cat. no. CRL-2522, RRID: CVCL_3653, U-2 OS, osteosarcoma, from ATCC cat. no. HTB-96, RRID: CVL_0042) were cultured in dedicated media supplemented with 10% fetal bovine serum (FBS) according to the ATCC instructions.

### Senescence induction

For DDR-sen, the cells were incubated with doxorubicin (Cayman Chemical, WI-38 and BJ 250 nM, U2OS 100 nM) for 24 h, washed and released into a fresh medium and cultured for additional days indicated in the figures. For Rep-sen, the cells were split every 3-4 d until the cells stopped proliferation with PDL >50 for WI-38 cells and >70 for BJ. For PMD-sen induction, cells were treated with the medium containing SDS (WI-38 0.009%, BJ 0.014%) for 24 h, washed twice with medium, covered with the fresh medium and incubated for additional days as indicated in the figures. The SDS concentration was optimized for each cell type and FBS lot. SDS failed to induce PMD-Sen in U2OS cells.

### SA-β-Gal assay

Cells were fixed for 3 min in 2% paraformaldehyde and incubated for approximately 16 h at 37°C with a staining solution containing 1 mg ml^−1^ X-gal, 40 mM citric acid in sodium phosphate buffer (pH 6.0), 5 mM K_4_[Fe(CN)_6_]3H_2_O, 5 mM K_3_[Fe(CN)_6_], 150 mM NaCl and 2 mM MgCl_2_. Additionally, nuclei were stained with DAPI.

### SA-β-Gal positive cells

SA-β-gal-positive cells were quantified using ImageJ (Fiji). For each image, a signal intensity threshold was established based on the SA-β-gal staining of the positive control sample to define positive staining. Cells exceeding this threshold in the cytoplasmic region were scored as SA-β-gal-positive. Total cell numbers were determined by counting DAPI-stained nuclei in the same field of view. The percentage of SA-β-gal-positive cells was calculated as the number of threshold-positive cells divided by the total number of DAPI-positive nuclei.

### siRNA transfection

Twenty-four hours after seeding, cells were transfected with siRNA using Lipofectamine RNAiMAX (Thermo Fisher Scientific) according to the manufacturer’s protocol. The total concentration of siRNA was 1 nM. Cells were analyzed at least after 72-96 h upon transfection and silencing efficiency was controlled by Western blotting or qPCR. Sequences of the used siRNAs: siPAK1#1 Seq1: GGATGGCTCTGTCAAGCTAATCTGAC, Seq2: GTCAGTTAGCTTGACAGAGAGCCATCCA, siPAK1#2 Seq1: CCCAAACATTTGTGAATTTACTTGAC, Seq2: GTCCAGTAATTCACAAATGTGTTTGGGT, siPAK1#3 Seq1: CTGTCCCTTGTTTATATAACATTGAGA, Seq2: CTCAAATGTTTATATACAAAGGGACAGGA, siPhafin2#1 Seq1: GTCATACTGCATTAGCTTCTACCTT, Seq2: AAGGTAGAAGCTAATGCAGTATGACAA, siRAC1#1 Seq1: GGTATTATCAGGAAATGTTTTTCTA, Seq2: TAAGAAAACATTTCCTGATATAATACCAA

### Immunofluorescence

For standard immunofluorescence, cells were seeded in full medium onto glass coverslips and incubated according to the experiment. At the end point, cells were fixed in 4% PFA for 10 min and treated with saponin 0.1% or Triton X-100 for permeabilization. Next, cells were washed twice with PBS for 5 min, blocked in Bullet One Histo for 30 min and incubated with appropriate primary antibodies for 1 h, followed by 30 min incubation with appropriate fluorescent secondary antibodies. DAPI and SiR-actin were used to stain nuclei and F-actin (for cell outline), respectively, and were added during incubation with fluorescent secondary antibodies. Cells were mounted with anti-fade mounting media.

### Dextran-uptake assay

Cells were incubated at 37°C with 70,000 MW dextran conjugated with different fluorescent dyes for 45-60 min (0.125 mg/ml) unless stated otherwise, washed with cold PBS five times, fixed and then the fluorescence and localization of internalized dextran was assessed by confocal microscopy. Where indicated, inhibitors were present in the medium for the entire duration of dextran incubation and uptake experiments.

### Dextran quantification

Dextran uptake was quantified using ImageJ. Fluorescence images were converted to 16-bit format, and an identical threshold was applied across all images within each experiment. Dextran-positive vesicles were detected using the Analyze Particles function with predefined minimum and maximum size parameters to exclude background and non-vesicular signals. The number of vesicles per image was normalized to the number of cells in the same field, and results are reported as number of macropinosomes per cell.

### WGA labeling

Cells were incubated with Alexa Fluor 488-conjugated wheat germ agglutinin (WGA; at 37 °C for 1 hour). After labeling, cells with or without concurrent dextran treatment were washed with PBS and fixed with 4% paraformaldehyde for subsequent immunostaining.

### Western blotting

Immunoblotting was performed following a standard protocol. Antibodies used in this study are listed in Table 1, and antibody validation was performed by the distributors. Images of blotted membranes were obtained by ChemiDoc Touch MP (Bio-Rad).

### Manual vacuole scoring

Vacuoles were scored manually by visually inspecting phase-contrast or holotomographic images. Cells containing one or more cytoplasmic vacuoles exceeding a predefined size threshold were classified as vacuole-positive. Size thresholds and scoring criteria were established based on control samples. Scoring was performed blinded to treatment groups to minimize bias.

### Nuclear area quantification

Nuclei were identified by DAPI staining and segmented using automated thresholding and watermark function in ImageJ (Fiji). The area of each nucleus was measured in pixels and converted to µm² using microscope calibration data.

### Confocal microscopy

Images were acquired using LSM 780, 880 or 900 confocal microscopes (Zeiss). At least seven 16-bit images with resolution of at least 2048×2048 pixels were acquired for each sample/experiment using the confocal microscopes with 40x/1.40 Oil Plan-Apo, 63x/1.46 α-Plan-Apo objectives. ZEN software (Zeiss) was used for acquisition. Signal quantification was performed using Fiji/ImageJ software. For time-lapse super-resolution imaging, a Nikon NSPARC microscope equipped with a 60×/1.40 Oil Plan Apo λ objective (WD 0.13) was used. Images were acquired using NIS-Elements software (version 5) and processed by Richardson-Lucy deconvolution.

### Holotomography

Holotomography is a transmitted-light microscopy technique that uses patterned illumination to acquire multiple images of a sample, which are then processed to reconstruct three-dimensional refractive index (RI) maps. An in-situ incubator was used for keeping the cells alive during the experiment. Cells were observed using 60× NA 1.2 water immersion objective lense installed on HT-X1 instrument. The hologram acquisition was captured as individual holographic images by scanning object illuminating beam. For time-lapse acquisition the images were captured on average every 7-45 seconds. The 3D distribution of the refractive index of live cells and fluorescence of FITC-dextran and their 2D maximum intensity projections were generated, visualized, and analyzed with TomoStudio software (Tomocube Inc, Korea). For the morphological analysis of the cells, TomoAnalysis software was used.

For holotomography coupled with fluorescence imaging of cells loaded with FITC-conjugated dextran, correlative 3D fluorescence images were acquired using the HT-X1 instrument. The data was imaged and visualized using a commercial softwares TomoStudio and TomoAnalysis (Tomocube Inc., Korea).

### Vesicle quantification in tomograms

Tomographic reconstructions were analyzed using a custom pipeline implemented in TomoAnalysis software (Tomocube Inc, Korea). For each dataset, only selected regions of interest (ROIs) containing individual cells were processed. Background suppression was performed by generating a 3D median filter (Median XYZ), followed by iterative arithmetic operations to merge the background model with the original tomogram and enhance vesicle contrast. Candidate vesicles were defined as round dark objects using a combination of 3D TopHat filtering and eigenvector analysis of the Hessian matrix to capture both large and small spherical structures. Detection was refined by global and local thresholding of refractive index (RI), including 3D RI Tophat correction. The segmented vesicles were merged through logical operations and labeled as distinct objects, with size filtering to exclude noise. Quantification was performed in three dimensions, yielding the vesicle number, area, and volume per cell within each ROI. All analyses were performed using identical filtering parameters and threshold settings across samples to ensure consistency and comparability.

### RNA extraction and qRT-PCR

qRT-PCR was performed following a protocol described by Suda et al.^7^. Cells were lysed in TRIzol. RNA was extracted and purified using Direct-zol RNA MicroPrep (Zymo Research) following the manufacturer’s instructions. Complementary DNA (cDNA) was synthesized using SuperScript IV VILO Master Mix (Thermo Fisher Scientific). The expression of target genes was determined using QuantStudio 1 Real-Time PCR system (Thermo Fisher Scientific). PCR was performed using PowerTrack SYBR Green Master Mix (Thermo Fisher Scientific) with primer pairs listed in Table 2. Changes in gene expression, expressed as fold change, were calculated using the ΔΔCt method, where ACTB was used as a reference gene for normalizing the expression. The plots were generated to show the fold changes of each replicate (GraphPad Prism). Unpaired Student’s *t*-test was used to check the significant changes of gene expression between pairs.

### RNA sequencing and sample preparation

Total RNA was extracted from WI-38 human lung fibroblasts using TRIzol reagent (Thermo Fisher Scientific). Cells were seeded at 3.5 × 10⁵ cells per 6 cm dish in triplicates for each condition: proliferating (PDL 37-39) and replicative senescent (PDL >51), treated with either siSCRAMBLE or siPAK1#1 (1 nM, transfected with Lipofectamine RNAiMAX). Medium was replaced 24 hours post-transfection, and RNA was harvested 72 hours later. For each condition, one additional replicate was used to assess RNA concentration and quality via Qubit fluorometric quantification, NanoDrop spectrophotometry, and Agilent Bioanalyzer. Twelve samples in total were submitted for RNA sequencing, stored in 1.5 mL tubes containing 250-300 µL of TRIzol, and labeled according to experimental condition and replicate. mRNA was enriched using the NEBNext Poly(A) mRNA Magnetic Isolation Module (NEB, Cat# E7490), followed by strand-specific cDNA synthesis and library preparation using the NEBNext Ultra II Directional RNA Library Prep Kit for Illumina (NEB, Cat# E7760), according to the manufacturer’s protocol. Sequencing was performed on the Illumina NovaSeq X Plus platform using 1.5B paired-end flow cells (150 bp, 300 cycles). One sequencing lane was used for all 12 samples, with a target output of 40 million paired end reads per sample.

### RNA-seq data analysis

RNA-seq data were processed and analyzed using the usegalaxy.org platform with standard parameters unless otherwise noted. Raw sequencing reads (FASTQ files) were first evaluated for quality using FastQC. Reads were aligned to the *Homo sapiens* reference genome (hg38) using HISAT2. Aligned reads were quantified at the gene level using featureCounts, with GENCODE/Ensembl annotations. Two parallel approaches were used for differential gene expression analysis: (1) DESeq2, which models count data using a negative binomial distribution, and (2) Limma with voom transformation, which estimates mean-variance relationships from count data and applies linear modeling. Both methods used default settings with Benjamini-Hochberg correction for multiple testing. Genes with an adjusted p-value < 0.05 or p-value < 0.05 and log₂FoldChange > 1 or < 1 were considered differentially expressed.

Principal component analysis (PCA) and distance matrix were used to assess sample clustering and variance between groups. Heatmaps were generated for genes of interest using custom gene lists, with expression values scaled by row. Downstream analyses were performed in R. Functional enrichment analysis was performed using clusterProfiler for KEGG pathway terms. Significantly enriched terms were filtered using p-value threshold (< 0.05) of downregulated genes (log₂FoldChange < 1) and visualized with bar plots. Venn diagrams were used to identify overlapping or unique gene sets between comparisons.

### Stable protein expression by lentiviral transduction

Stable protein expression was achieved using a lentiviral transduction system produced in HEK293T cells. HEK293T cells were seeded in 6-well plates at a density of 3.0 × 10⁵ cells per well in 2 mL of DMEM supplemented with 10% FBS, L-glutamine, and penicillin/streptomycin. Cells were cultured overnight and transfected the following day once 80–90% confluence was reached. Lentiviral particles were generated by transient transfection using a three-plasmid system consisting of the lentiviral transfer vector, pCAG-HIVgp (Gag/Pol), and pCMV-VSVG-RSV-Rev (VSV-G and Rev), mixed at a ratio of 2:1:1. Polyethylenimine (PEI) was added at a DNA:PEI ratio of 1:2 (w/w). DNA–PEI complexes were incubated for 5-10 minutes at room temperature and then added dropwise to the cells. Transfections were performed in DMEM without FBS and allowed to proceed for 12-20 hours. After that time, medium was replaced with 1.1 mL of fresh medium appropriate for the destination cells, supplemented with forskolin at a final concentration of 10 µM to enhance viral production. Viral supernatants were collected after 24 hours of incubation. In parallel, destination cells were seeded in 6-well plates at a density of 1.0-2.0 × 10⁵ cells per well in 2 mL of complete growth medium to reach near confluence at the time of infection. For infection, viral supernatants were filtered through a 0.45 µm PVDF membrane and added directly to the destination cells. After 24 hours medium in destination cells was replaced to full DMEM. Selection of stably transduced cells was initiated approximately 48 hours post-infection by incubation in medium containing the appropriate selection antibiotic. Antibiotic concentrations were optimized per cell line. Control cells lacking resistance were plated in parallel to monitor selection efficiency. Antibiotic-resistant populations were expanded and maintained under selection. Phenotypic analyses, including imaging, were performed after stable expression was confirmed.

### Statistical analysis

Statistical analyses for all experimental data were performed using GraphPad Prism software. Data are presented as mean ± standard error of the mean (SEM), unless stated otherwise. Comparisons between two experimental groups were conducted using unpaired or paired two-tailed Student’s *t*-tests. A p-value of less than 0.05 was considered statistically significant. The number of biological replicates and exact statistical tests used are specified in the corresponding figure legends.

## Acknowledgements

We are grateful to M. Tanigawa and Y. Moriyama for critical reading of the manuscript. We are grateful to Y. Moriyama and R. Ishida for providing plasmids to study endosome populations. We thank K. Koizumi, S. Komoto, and P. Barzaghi for technical assistance with confocal microscopy. A portion of this work was performed with the help of OIST Imaging Section members. We thank the Tomocube team for holotomography demonstration and pipeline creation for quantification. We thank OIST Sequencing facility for performing RNA-seq. We acknowledge C. Kang, F. Meitinger, and Y. Lee for their valuable discussions and guidance on specific experiments.

## Author contributions

KKoz conceived the study, designed and performed the experiments, analyzed the data, and wrote the original manuscript draft. NR performed qPCR experiments and reviewed the manuscript. KKon contributed to methodology, participated in manuscript review and editing, supervised the study and secured funding. All authors reviewed and approved the final manuscript.

## Supplementary Tables

**Table 1.** Primary antibodies used for western blotting.

| Name | Species | Catalog number | RRID | MW (kDa) | Dilution used for WB |
| --- | --- | --- | --- | --- | --- |
| Lamin B1 | Rabbit | ab133741 | AB_2616597 | 66 | 1:1000 |
| p21 | Rabbit | ab109520 | AB_10860537 | 21 | 1:500 |
| p16 | Rabbit | ab108349 | AB_10858268 | 16 | 1:500 |
| α-tubulin | Rabbit | CS #2144 | AB_2210548 | 52 | 1:2500 |
| p53 | Mouse | sc-126 | AB_628082 | 53 | 1:500 |
| GAPDH | Rabbit | CS #5174 | AB_10622025 | 37 | 1:2500 |
| p-SMAD2 | Rabbit | CS 3108S | AB_490941 | 60 | 1:200 |
| p-SMAD3 | Rabbit | CS 9520S | AB_2193207 | 52 | 1:200 |
| β-actin | Rabbit | #12004164 | AB_2861334 | 42 | 1:1000 |
| PAK1 | Rabbit | CS 2602S | AB_330222 | 68 | 1:1000 |
| Cyclin B1 | Rabbit | CS 12231 | AB_2783553 | 55 | 1:500 |
| p-Histone H3 | Rabbit | ab1791 | AB_302613 | 17 | 1:1000 |
| Histone H3 | Rabbit | CS 53348 | AB_2799431 | 17 | 1:1000 |

**Table 2.**
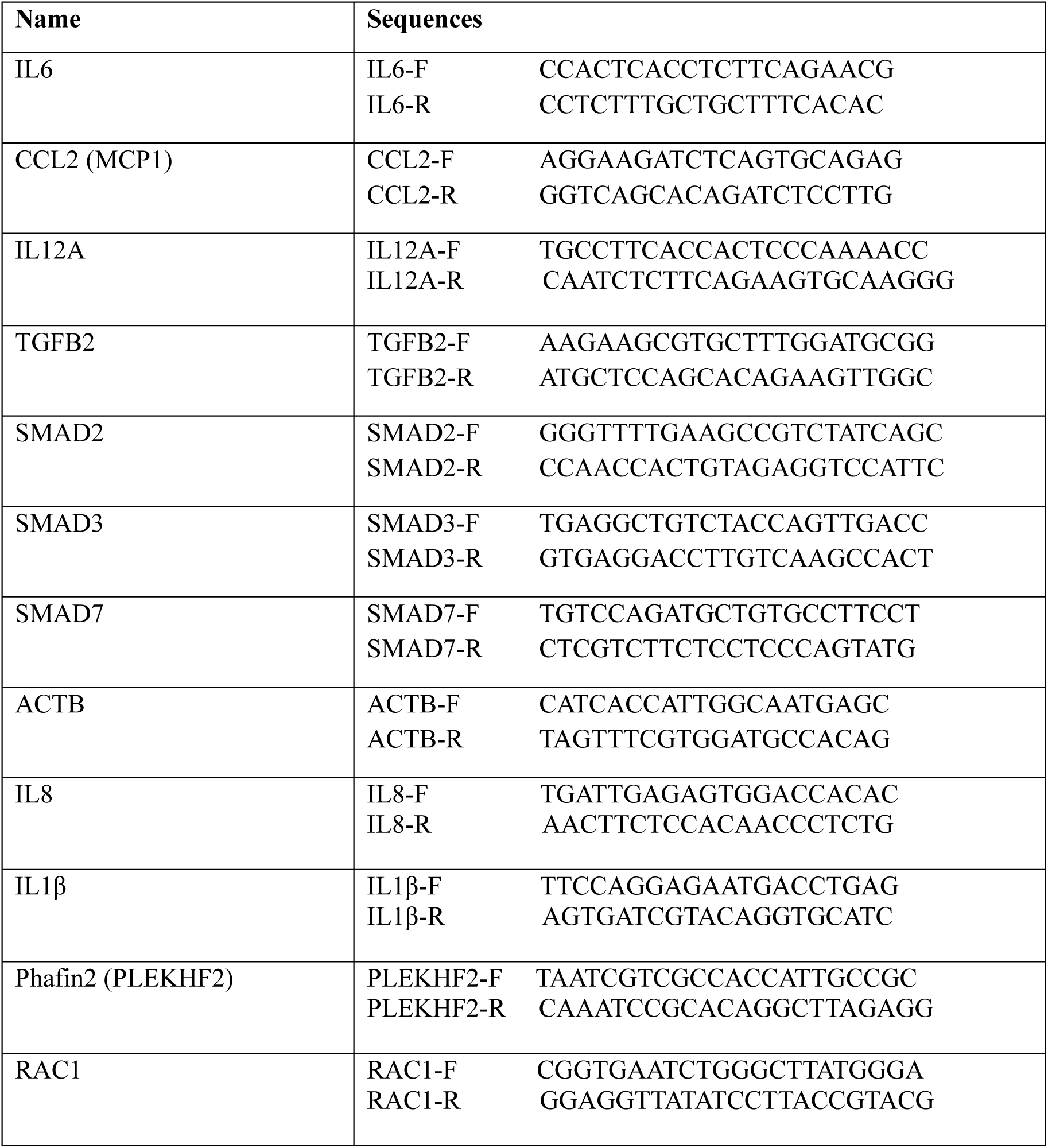
Primers used for qRT-PCR.

**Table 3.** List of inhibitors of activators.

| Function | Target | Name | Cat. Number | Final concentration |
| --- | --- | --- | --- | --- |
| Cytoskeleton | Actin | Latrunculin A | 125-0436 | 200nM |
| Cytoskeleton | Actin | Cytochalasin D | 11330 | 500nM |
| PAK1 inhibitor | PAK1 | NVS-PAK-1 | HY-100519 | 100nM |
| Macropinocytosis inhibitor | PI3K | LY294002 | HY-10108 | 1μM |
| Macropinocytosis inhibitor | macropinocytosis | Imipramine | HY-B1490 | 1μM |
| Macropinocytosis inhibitor | Na<sup>+</sup>/H<sup>+</sup>exchanger (NHE) inhibitor | EIPA | HY101840A | 10μM (maximum sublethal) |
| Macropinocytosis inhibitor | PAK1 | IPA-3 | HY-15663 | 1μM |
| PAK1 activator | PAK1 | Fingolimod | HY-12005 | 5μM |
| TGFβ activator | TGFβ | Recombinant human TGFB1 | 75362S | 5ng/mL |

**Table 4.** Fluorescent reagents.

| Target | Name | Cat. No. | Excitation | Emission | Concentration |
| --- | --- | --- | --- | --- | --- |
| Membranes | WGA | W11261 | 495 | 519 | 5μg/ml |
| Macropinocytosis cargo | Dextran (70.000 MW) | D1864 | 595 | 615 | 0.125 mg/mL |
| Macropinocytosis cargo | Dextran (70.000 MW) | 46945 | 492 | 518 | 0.125mg/mL |
| Growth factor | EGF | E13345 | 495 | 519 | 100ng/mL |

**Table 5.**
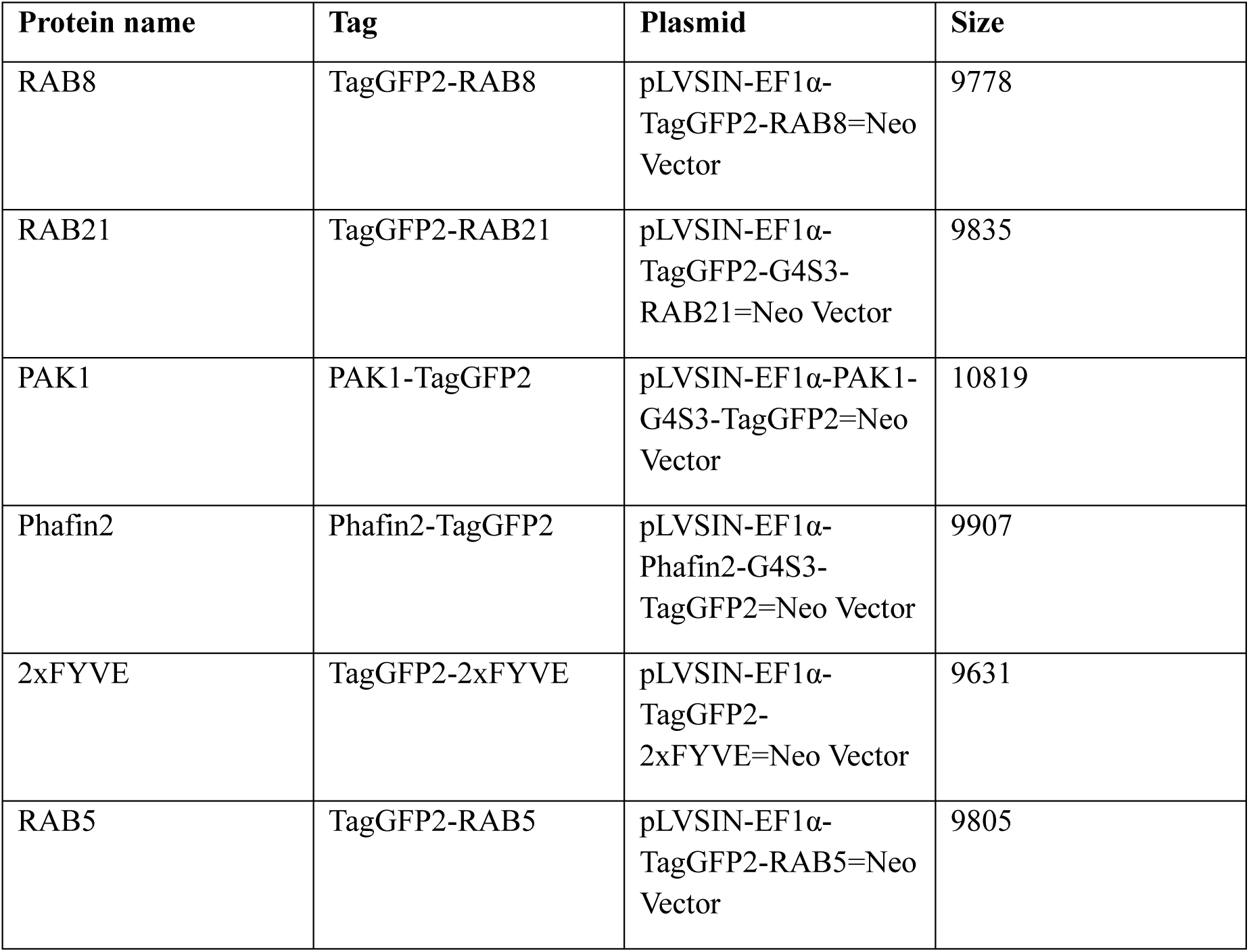
Plasmids used for expression of endosomal markers in U2OS cells.

