## Supplementary Figures for "Constitutive macropinocytosis sustains the inflammatory phenotype of senescent cells"

### Supplementary Figures and Movies and their legends

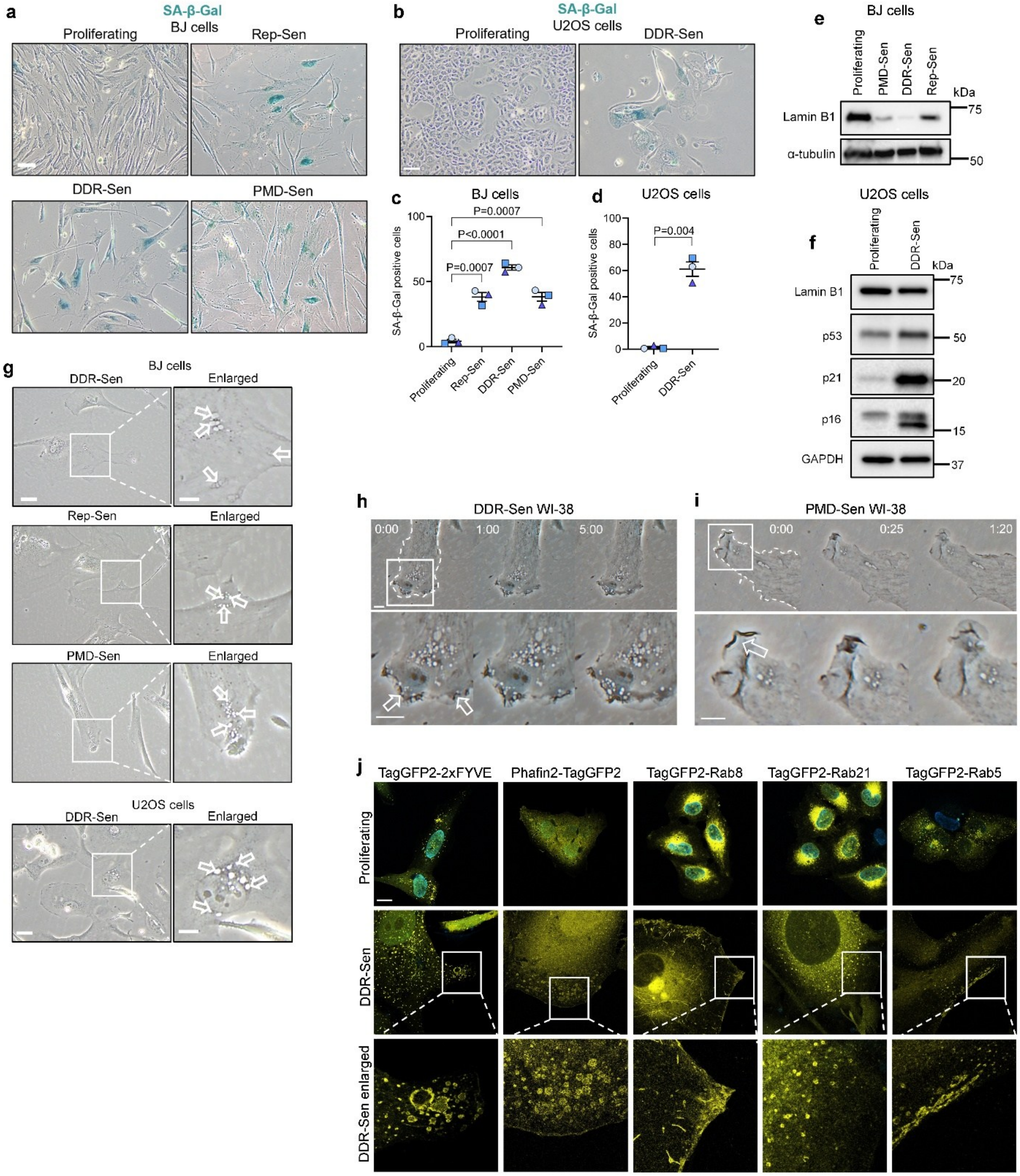


**Supplementary Figure 1. Senescent cells form a high number of peripheral cytoplasmic large vesicles. (a, b)** SA-β-Gal positive cells detected after replication arrest (Rep-Sen) or 16 days after treatment (doxorubicin – DDR-Sen, SDS –

PMD-Sen) in BJ (a) and U2OS (b) cells. Scale bar 20 µm. **(c, d)** Quantification of A. Approximately 50 cells were counted per experiment. n=3. **(e, f)** Western blotting using lysates of proliferating and senescent BJ or U2OS cells. The lysates were collected 16 days after treatment with SDS or doxorubicin. **(g)** Phase-contrast analysis of vacuole number in senescent BJ or U2OS cells. Scale bar 10 µm. Arrows mark the phase-bright cytoplasmic vacuoles. **(h)** Representative frames from time lapse recordings (Movie 2 and 3) of vesicles formation in DDR-Sen and PMD-Sen WI-38 cells. Scale bar 10 µm. Time format m:ss. **(j)** Representative fluorescence images of proliferating and doxorubicin-induced senescent U2OS cells stably expressing GFP-tagged endosomal markers: FYVE (PtdIns3P), Phafin2, Rab5 (early endosomes), Rab8, and Rab21 (macropinosome-associated Rab GTPases). GFP-tagged proteins are shown in yellow; nuclei were counterstained with DAPI (cyan). Scale bar 20 µm.

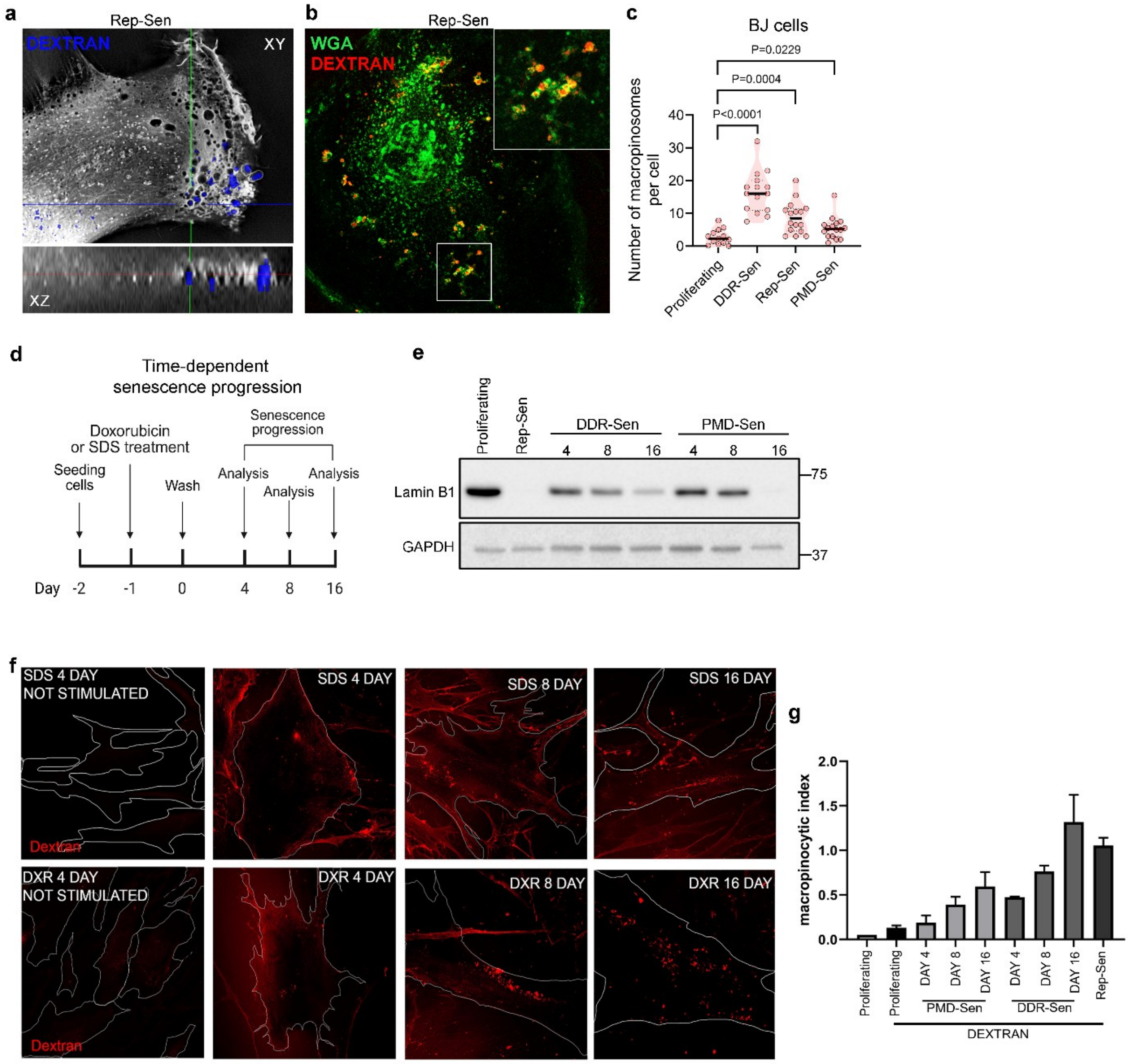


**Supplementary Figure 2. Senescent cells upregulate macropinocytosis. (a)** Holotomography analysis of fluorescent dextran (blue) localization in Rep-Sen WI-38. **(b)** Confocal microcopy analysis of fluorescent dextran (red) vesicular localization (WGA-green) in Rep-Sen WI-38**. (c)** Dextran uptake in BJ proliferating and senescent cells. **(d)** Scheme of senescence progression and sample analysis time points **(e)** Western blotting using lysates of proliferating and senescent WI-38 cells. The lysates were collected 4, 8 and 16 days after treatment with SDS or doxorubicin and after induction of replicative senescence. **(f)** Dextran uptake in PMD-Sen (SDS) and DDR-Sen (DXR) WI-38 cells on day 4, 8 and 16 after senescence induction. Not stimulated – no dextran incubation. **(g)** Quantification of macropinocytosis index (cell size / macropinosome number) in proliferating and senescent WI-38 cells at different time points of senescence progression. n=3.

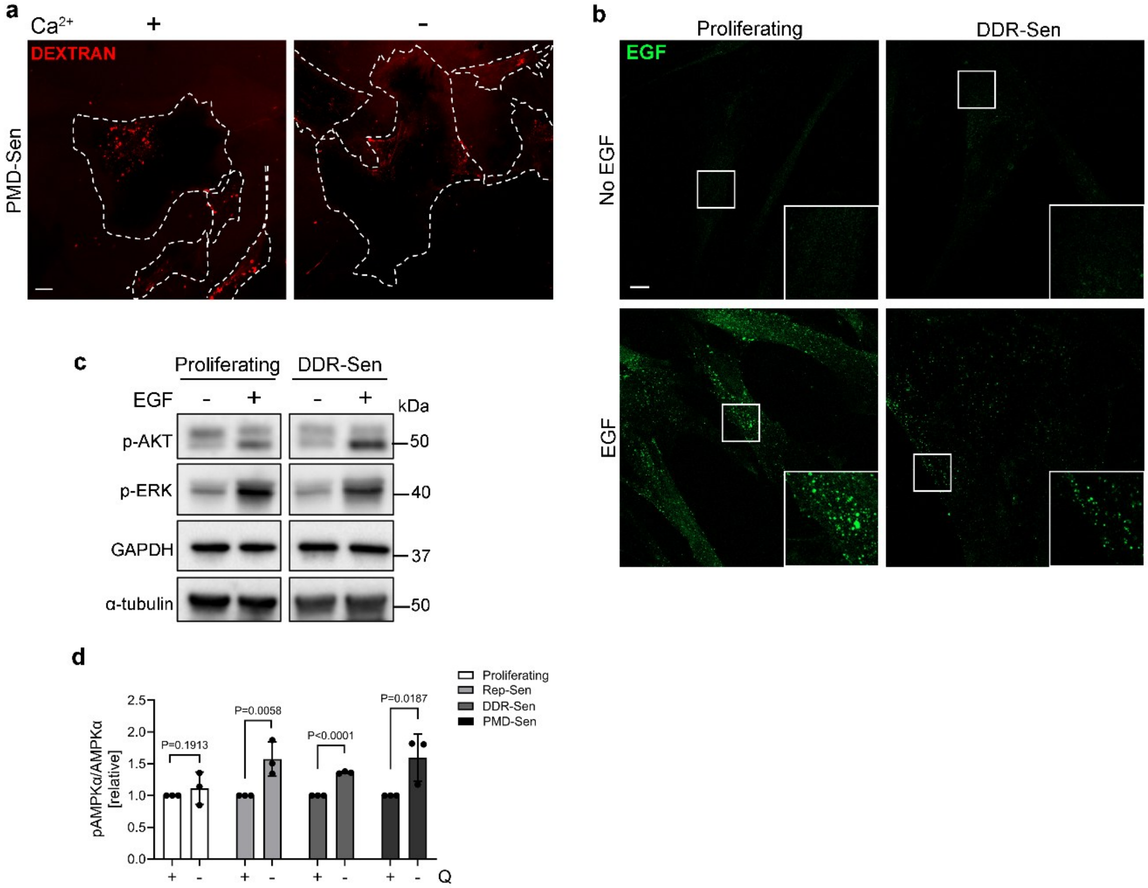


**Supplementary Figure 3. Macropinocytosis in senescent cells is glutamine and $Ca^{2+}$ sensitive and EGF-independent. (a)** Confocal microscopy analysis of fluorescent dextran uptake in PMD-Sen WI-38 cells. Cells were incubated in medium without extracellular calcium 30 min before dextran loading. Scale bar 20 µm. **(b)** Confocal microscopy analysis of fluorescent EGF (green) endocytosis in proliferating and senescent WI-38 cells. Cells were incubated in medium with or without EGF for 60 min. Scale bar 20 µm. **(c)** Western blotting using lysates of proliferating and senescent WI-38 cells. The lysates were collected 16 days after treatment with doxorubicin and after 15 min incubation with EGF. **(d)** Quantification of immunoblots in Fig. 4f. WI-38 proliferating and senescent cells were incubated in medium with or without glutamine for 24 hours. n=3.

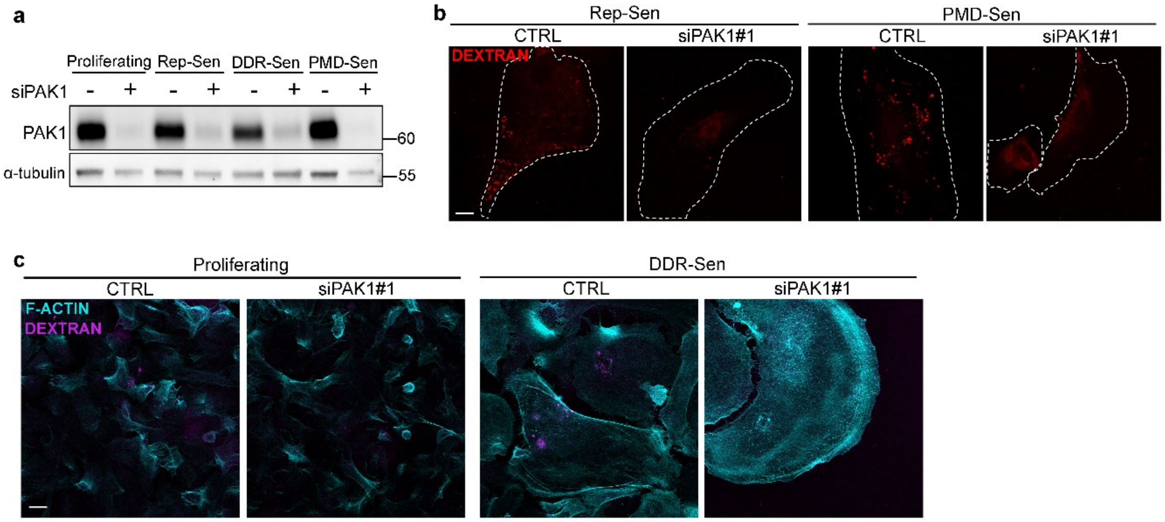


**Supplementary Figure 4. PAK1 is required for macropinocytosis in senescent cells. (a)** Efficiency of siRNA-mediated knockdown of PAK1. **(b)** Confocal microscopy analysis of fluorescent dextran (red) uptake in Rep-Sen and PMD-Sen WI-38 cells upon PAK1 knock down. Scale bar 20 µm. **(c)** Confocal microscopy analysis of fluorescent dextran (magenta) uptake in proliferating and senescent U2OS cells upon PAK1 knockdown. F-actin (cyan) was stained with siR-Actin for 30 min. Scale bar 20 µm.

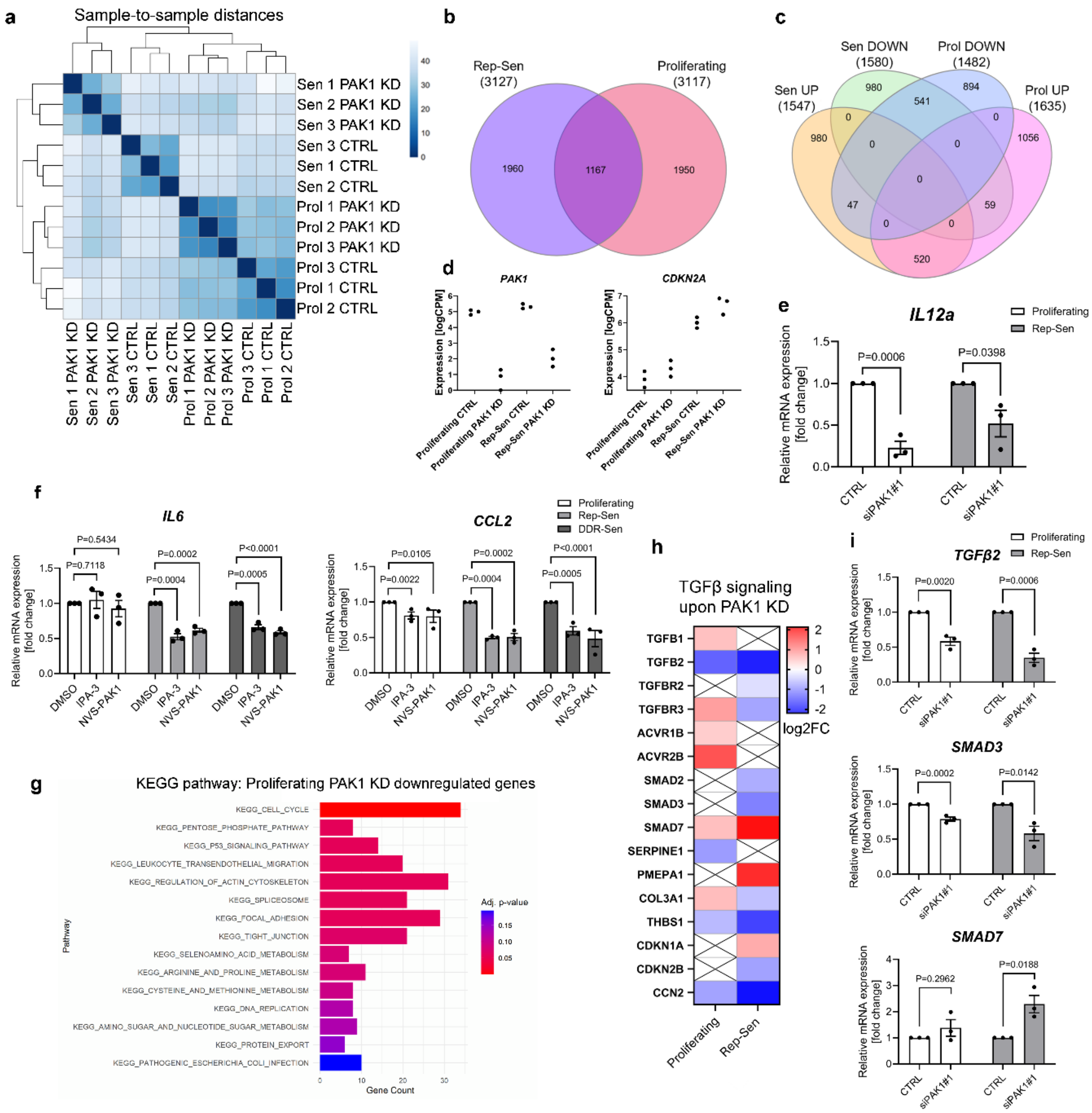


**Supplementary Figure 5. PAK1 knockdown downregulates IL6, CCL2 and IL12a expression in senescent cells. (a)** Sample-to-sample distance matrix showing clustering of RNA-seq replicates across four groups: proliferating control (Prol CTRL), proliferating PAK1 knockdown (Prol PAK1 KD), replicative senescent control (Sen CTRL), and replicative senescent PAK1 knockdown (Sen PAK1 KD) **(b)** Venn diagram illustrating the overlap of differentially expressed genes (DEGs) between replicative senescent and proliferating WI-38 cells following PAK1 knockdown. **(c)** Venn diagram showing the overlap of significantly upregulated (UP; log2FC > 1) and downregulated (DOWN; log2FC < −1) DEGs between replicative senescent and proliferating WI-38 cells upon PAK1 knockdown. **(d)** RNA-seq expression levels of selected genes, PAK1 and CDKN2A, across all conditions, presented as logCPM (counts per million), validating knockdown efficiency and senescence-associated gene expression. **(e)** qPCR analysis of IL12A expression in proliferating and replicative senescent (Rep-Sen) WI-38 cells following PAK1 knockdown. n=3 **(f)** qPCR analysis of IL6 and CCL2 expression in proliferating, replicative senescent (Rep-Sen), and DNA damage–induced senescent (DDR-Sen) WI-38 cells following pharmacological inhibition of PAK1 using IPA-3 and NVS-PAK1-1 for 72 h**.** n=3. **(g)** KEGG pathway enrichment analysis of significantly downregulated genes ($\log2FC < -1$, p-value $< 0.05$) in proliferating WI-38 cells following PAK1 knockdown. **(h)** Heatmap showing RNA-seq–derived expression changes of TGFβ pathway–associated genes in proliferating and replicative senescent WI-38 cells following PAK1 knockdown. **(i)** qPCR validation of RNA-seq findings demonstrating expression changes in TGFB2, SMAD3, and SMAD7 in proliferating and replicative senescent WI-38 cells upon PAK1 depletion. n=3.

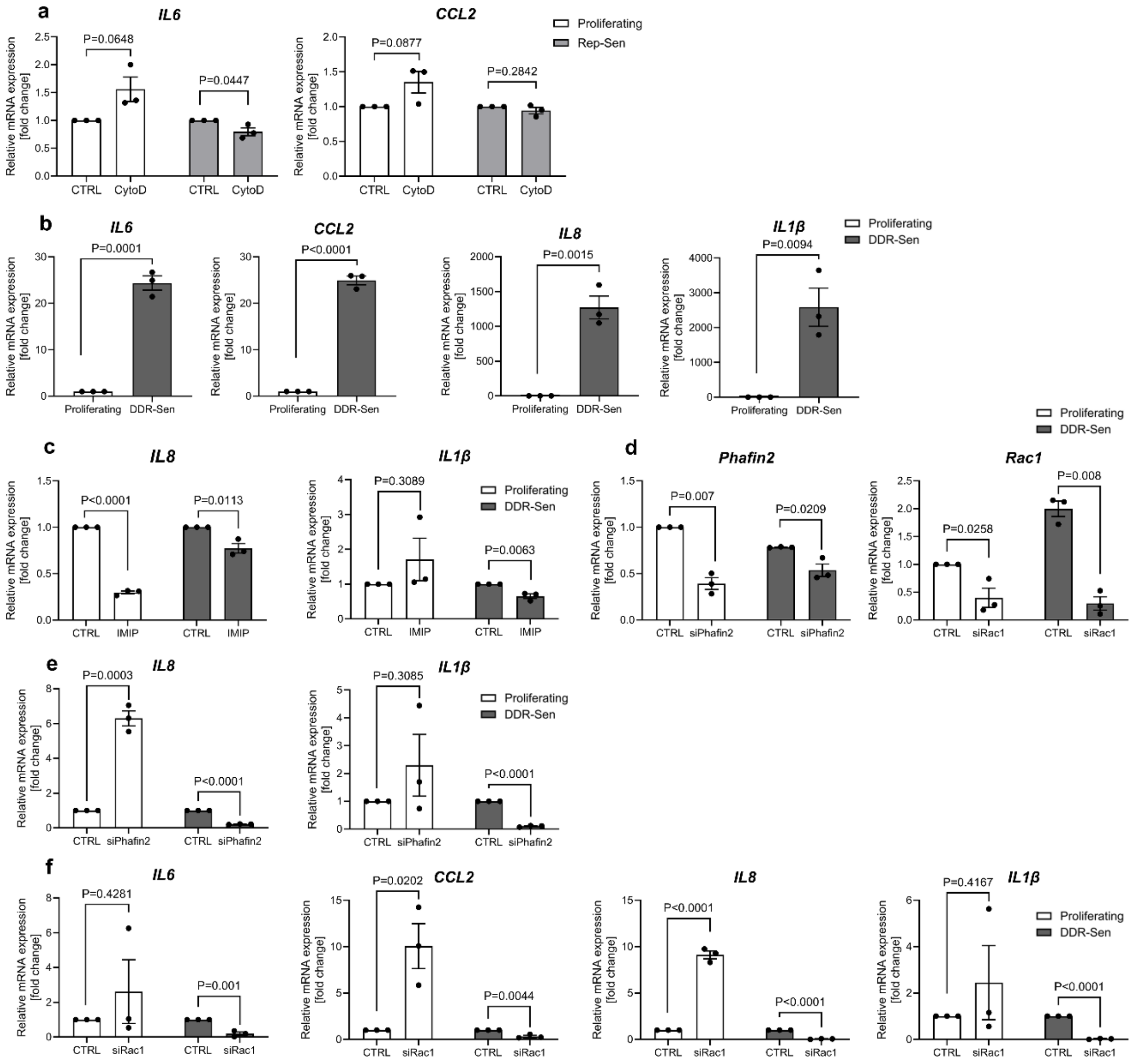


**Supplementary Figure 6. Macropinocytosis contributes to SASP expression. (a)** qPCR analysis of *IL6* or *CCL2* expression following treatment with Cytochalasin D (CytoD) in proliferating and Rep-Sen WI-38 cells. n=3 **(b)** qPCR analysis of *IL6*, *CCL2, IL8* and *IL1β* expression in proliferating and DDR-Sen U2OS cells. n=3 **(c)** qPCR analysis of *IL8* or *IL1β* expression following treatment with imipramine in proliferating and DDR-Sen U2OS cells. n=3. **(d)** qPCR analysis of *Phafin2* or *Rac1* expression in proliferating and DDR-Sen U2OS cells following control or Phafin2- or Rac1-targeting siRNA treatment. n=3 **(e)** qPCR analysis of *IL8* or *IL1β* expression following treatment with siRNA targeting Phafin2 in proliferating and DDR-Sen U2OS cells. n=3. **(f)** qPCR analysis of *IL6, CCL2, IL8* or *IL1β* expression following treatment with siRNA targeting Rac1 in proliferating and DDR-Sen U2OS cells. n=3.

#### Movie legends

**Movie 1-4. Time-lapse of peripheral large vesicles formation in WI-38 cells.** WI-38 cells were monitored by phase-contrast microscopy for 5-15 minutes. Movie speed was increased 16 times.
Movie 1 - Rep-Sen WI-38, Movie 2 -DDR-Sen WI-38, Movie 3 - PMD-Sen WI-38, Movie 4 - Proliferating WI-38.

**Movie 5-8. Dynamics of large vesicle formation and maturation in senescent WI-38 cells by holotomography.** WI-38 cells were monitored by holotomography microscopy for 10-90 minutes. Images were acquired every 7-45 s., Movie 5 - DDR-Sen WI-38, Movie 6 - PMD-Sen WI-38, Movie 7 - Proliferating WI-38, Movie 8 - DDR-Sen WI-38. Scale bars and time are indicated in the movies.

**Movie 9-10 Dynamics of Phafin2-GFP and GFP-2xFYVE vesicle formation and maturation in senescent U2OS cells.** U2OS senescent cells stably expressing 2xFYVE-GFP or Phafin2-GFP were monitored by super-resolution Nikon NSPARC microscopy for the time indicated in the movies. Images were acquired every 3s. Scale bars and time are indicated in the movies.
